# Neuroprotective function of astrocyte p75^NTR^ in Alzheimer’s Disease through regulation of cholesterol metabolism

**DOI:** 10.1101/2025.10.20.683521

**Authors:** Xueyan Han, Meng Xie, Carlos F. Ibáñez

**Affiliations:** College of Biological Sciences, China Agricultural University, 100193 Beijing, China; Chinese Institute for Brain Research, Zhongguancun Life Science Park, 102206 Beijing, China; Peking University School of Psychological and Cognitive Sciences, 100871 Beijing, China; Department of Medicine Huddinge, Karolinska Institute, Huddinge 14157, Sweden; Peking University School of Life Sciences, PKU-IDG/McGovern Institute for Brain Research, Peking-Tsinghua Center for Life Sciences, 100871 Beijing, China; Department of Neuroscience, Karolinska Institute, Stockholm 17177, Sweden; Stellenbosch Institute for Advanced Study, Wallenberg Research Centre at Stellenbosch University, Stellenbosch 7600, South Africa

**Author notes:** Corresponding author: C.F.I.

**Keywords:** Aβ, amyloid precursor protein, neurotrophin, SREBP2, statins, 5xFAD

## Abstract

Reactive astrogliosis in Alzheimer’s Disease (AD) involves profound changes in the morphology, metabolism and secretion profile of astrocytes, but whether astrogliosis is beneficial or harmful, and under which conditions, remain open questions. Here we report an unexpected neuroprotective function of death receptor p75^NTR^ in astrocytes through its ability to regulate cholesterol metabolism. AD knock-in mice expressing signaling-deficient p75^NTR^ variants in astrocytes showed enhanced Aβ burden, brain histopathology and cognitive impairment, even when variants were introduced late in the disease process. Astrocytes expressing dysfunctional p75^NTR^ variants showed impaired uptake of Aβ oligomers, and their conditioned medium enhanced Aβ production in AD neurons. p75^NTR^ signaling negatively regulated astrocyte cholesterol biosynthesis and secretion, while cholesterol depletion restored Aβ uptake in mutant astrocytes and reduced Aβ production in AD neurons. In agreement with the role of astrocyte-derived cholesterol, statin treatment reverted the effects of astrocyte p75^NTR^ mutants on AD neuropathology. Thus, although neuronal p75^NTR^ has been widely recognized to amplify AD, astrocyte p75^NTR^ plays a neuroprotective role.

## Introduction

The abnormal processing of amyloid precursor protein (APP), aggregation of Aβ, Tau hyper-phosphorylation and propagation of these abnormally folded proteins have been collectively recognized as hallmarks of the initial, “biochemical” phase of Alzheimer’s Disease (AD) ^1^. The proteonopathy of the biochemical phase is thought to lead to a “cellular” phase of AD involving extensive feedback and feedforward interactions between multiple cell subpopulations, including not only neurons but also glial and vascular cells ^1^. Among these, astrocytes are central players in the cellular phase of AD. Astrocytes are one of the most abundant cell types in the brain and perform several critical functions, including maintenance of the blood–brain-barrier (BBB), neurotransmitter recycling, regulation of synaptic plasticity and brain metabolism, as well as clearing and degradation of Aβ ^2,3^, all of which are dysfunctional in AD. In response to injury and brain damage, astrocytes undergo pronounced morphological and transcriptional changes collectively known as astrogliosis ^4^. In AD, astrogliosis has been linked to both beneficial and harmful effects depending on the stage of disease progression as well as the brain region affected. To date, however, the molecular and cellular aspects of astrocyte reactivity in AD, as well as their precise contribution to pathogenesis and disease progression, remain poorly understood.

The p75 neurotrophin receptor (p75^NTR^), a member of the “death” receptor superfamily, can directly interact with APP ^5,6^ and is widely recognized as an important contributor to AD-associated neuropathology by accelerating Aβ production, neurite degeneration and cognitive impairment in mouse models of AD ^6–8^. Small molecule modulators of p75^NTR^ activity are currently being tested in clinical trials on AD patients ^9^. Death receptors, which also include the tumor necrosis factor receptor 1 (TNFR1), CD40, Fas, and others, induce cell death pathways as a mechanism for clearing damage produced after a lesion or insult. However, following severe injury or disease, they can also amplify tissue damage as a result of overactivation and/or overexpression Ibanez 2012. Indeed, expression of p75^NTR^ is increased in the brain of AD patients ^10–13^ as well as animal models of AD ^6,7,14^. p75^NTR^ functions as a receptor for neurotrophins, a family of neurotrophic growth factors that includes nerve growth factor (NGF) and brain-derived neurotrophic factor (BDNF), as well as other, structurally unrelated ligands, including myelin-derived components and the Aβ peptide (for review, see ^15,16^). Some of the main downstream signaling pathways engaged by p75^NTR^ include the NF-kB, RhoA and c-Jun Kinase (JNK) pathways ^17–21^. The neurotrophic functions of the neurotrophins are mainly mediated by the Trk family of receptor tyrosine kinases ^22,23^, which have the ability to overcome p75^NTR^ cell death signaling when co-expressed in the same cells, such as for example forebrain cholinergic neurons ^24^. In the healthy adult brain, p75^NTR^ is normally expressed at low levels in subpopulations of neurons and glial cells, including oligodendrocytes and astrocytes. In the latter, p75^NTR^ can be highly induced upon a variety of insults, including seizures, ischemia and neuroinflammation ^25,26^. A more recent study also reported induction of astrocyte p75^NTR^ expression in a mouse model of AD ^27^. Interestingly, under several of these conditions, astrocytes also upregulate expression of p75^NTR^ ligands, particularly pro-neurotrophins such as proNGF, which can interact strongly with p75^NTR^ but not with Trk receptors^24,28^.

In a previous study, we reported that signaling-deficient variants of p75^NTR^ conferred greater neuroprotection than the removal of the receptor (i.e. knock-out) when introduced in the endogenous *Ngfr* locus (which encodes p75^NTR^) of the 5xFAD mouse model of AD ^6^. These variants included a deletion of the receptor death domain (herein referred to as ΔDD) and the replacement of transmembrane residue Cys^259^ for Alanine (herein termed C259A). Although still competent to bind neurotrophins, neither ΔDD nor C259A p75^NTR^ molecules are able to signal in response to these ligands ^29,30^. Mechanistically, the beneficial effects of these variants on AD progression was explained by their ability to interfere with the internalization and intracellular trafficking of APP in neurons, leading to enhanced APP cleavage by plasma membrane alpha-proteases, reduced intracellular APP co-localization with BACE, and lower levels of Aβ species and amyloid plaque burden in the brain of 5xFAD mice ^6^. As these variants were introduced constitutively in all cells from birth, it remained unclear what effects, if any, they may have in other brain cells besides neurons, which are the major cell type producing Aβ in AD. In the present study, we generated conditional mouse *Ngfr* alleles for both the ΔDD and C259A p75^NTR^ variants and investigated their effects on AD progression when specifically introduced in astrocytes. Based on what was known until now about p75^NTR^ and its function in AD and other conditions, our expectation was that signaling-deficient variants of this receptor would also confer beneficial effects when expressed in astrocytes. Unexpectedly, we found that the opposite was the case.

## Results

### Accelerated and enhanced Aβ burden in the brain of 5xFAD mice expressing signaling-deficient p75^NTR^ variants in astrocytes

In order to test the functionality of signaling-deficient p75^NTR^ variants in a cell type-specific and time-dependent manner, we generated conditional alleles of the mouse *Ngfr* gene encoding p75^NTR^ (herein termed *ΔDD^fl^* and *C259A^fl^*) that allow expression of mutant ΔDD and C259A variants after Cre-mediated recombination (Supplementary Figure S1A, B). We verified that: i) the engineered alleles express normal levels of p75^NTR^ prior to recombination (Supplementary Figure S2A, B), and ii) mutant protein was expressed at normal levels in astrocytes after Cre-mediated recombination (Supplementary Figure 2C, D). In agreement with previous studies ^27^, we observed upregulation of p75^NTR^ in hippocampal astrocytes of the 5xFAD mouse model of AD (Supplementary Figure 3A, B). 5xFAD mice show a progressive increase in Aβ plaque burden in several brain regions, including hippocampus, cortex and thalamus, from 4 months onwards (Figure 1A, B). To study the role of astrocyte p75^NTR^ in AD pathogenesis, we utilized the *Aldh1l1-Cre^ER^*^T2^ driver which induces robust and specific recombination in astrocytes after tamoxifen treatment ^31^. Mice were generated carrying the *Aldh1l1-Cre^ER^*^T2^ driver, the 5xFAD transgene, and the *DD^fl/fl^* and *C259A^fl/fl^* conditional alleles. Tamoxifen was injected at 2 months of age to mutant and control 5xFAD mice to induce recombination prior to the overt manifestation of AD pathology, while avoiding possible developmental effects. Unexpectedly, histological analysis of brains at 4, 6 and 9 months revealed increased Aβ accumulation in hippocampus, cerebral cortex and thalamus of p75^NTR^ mutant 5xFAD mice compared to 5xFAD controls (Figure 1A, B). Subsequently, we administered tamoxifen at 6 months and examined brains at 9 months to assess whether signaling-deficient p75^NTR^ variants can also affect AD progression when expressed in astrocytes during an already established disease process. We found again significantly increased Aβ burden in the hippocampus, cortex and thalamus of p75^NTR^ mutants compared to controls (Figure 1A, B). These observations suggest that normal p75^NTR^ signaling in astrocytes contributes to restrain the development of Aβ aggregates in the AD brain both at the onset of and throughout the disease process.

**Figure 1.**
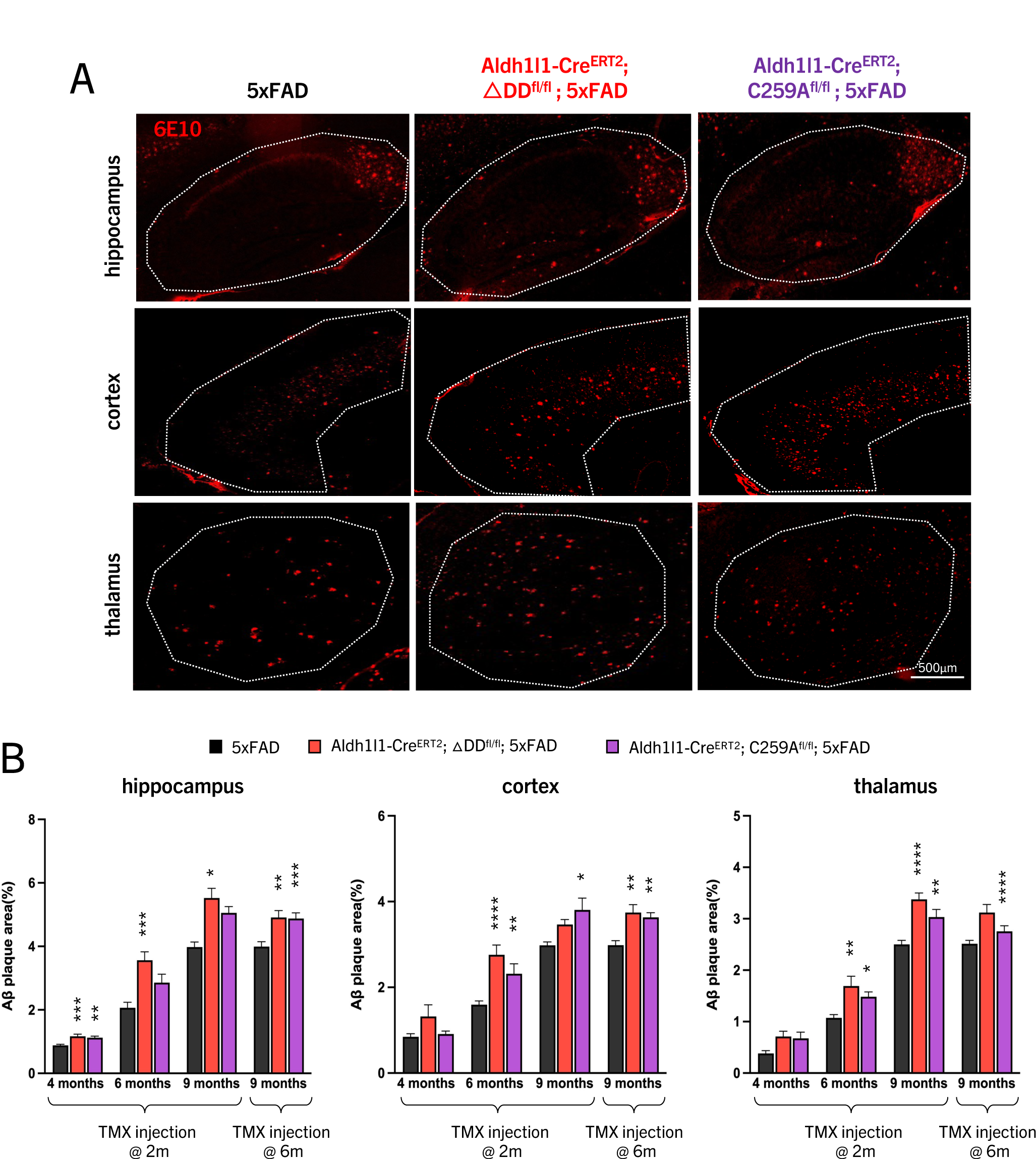
Accelerated and enhanced Aβ burden in the brain of 5xFAD mice expressing signaling-deficient p75^NTR^ variants in astrocytes. (A) Representative micrographs showing Aβ plaque depositions (as revealed by immunohistochemistry with 6E10 antibody) in sagittal sections through the hippocampus, cerebral cortex and thalamus of 5xFAD, Aldh1l1-CreER^T2^;ΔDD^fl/fl^;5xFAD and Aldh1l1-CreER^T2^;C259A^fl/fl^;5xFAD mice at 6 month of age. Mice were injected with tamoxifen at 2 month of age. The areas used for quantification are outlined. (B) Quantification of Aβ plaque area (as % of outlined region) in hippocampus, cerebral cortex and thalamus of 5xFAD, Aldh1l1-CreER^T2^;ΔDD^fl/fl^;5xFAD and Aldh1l1-CreER^T2^;C259A^fl/fl^;5xFAD mice injected with tamoxifen (TMX) at 2 month or 6 month of age and analyzed at 4, 6 or 9 month as indicated. Results are presented as mean ± SEM (N=7-11 mice per group) and analyzed by two-way ANOVA followed by Tukey’s multiple comparisons test. *, p<0.05; **, p<0.01; ***, p<0.001; ****, p<0.0001 vs. 5xFAD of the corresponding age and condition.

### Increased brain AD histopathology and learning/memory deficits in 5xFAD mice expressing signaling-deficient p75^NTR^ variants in astrocytes

5xFAD mice develop progressive astrogliosis and microgliosis in hippocampus, cerebral cortex and thalamus at 4, 6 and 9 months of age. In line with their increased Aβ burden, the brains of 5xFAD mice expressing signaling-deficient p75^NTR^ variants in astrocytes starting at 2 months of age showed enhanced gliosis compared to 5xFAD mice expressing the wild type receptor at the three ages tested (Figure 2A-D). With regards to astrogliosis, the enhancement produced by the mutant p75^NTR^ variants was even more pronounced among the disease-associated subpopulation, marked by expression of Vimentin ^32^ (Figure 2E, F). Even when mutant p75^NTR^ expression was initiated by tamoxifen injection at 6 months, did the brains of those mice also show enhanced microgliosis and Vimentin-associated astrogliosis at 9 months (Figure 2B, D, F), indicating the continued importance of astrocyte p75^NTR^ activity during the disease process. Neurite dystrophy in dendritic arbors of hippocampal and cortical neurons was as assessed by accumulation of reticulon 3 (RTN3), a protein known to form aggregates concentrated in and around Aβ plaques in the brains of both AD patients and APP transgenic mice ^33^. Here again, expression of mutant p75^NTR^ variants in astrocytes amplified RNT3-associated neurite dystrophy (Figure 2G, H). Together, these results show that increased Aβ burden in 5xFAD mice expressing signaling-deficient p75^NTR^ variants in astrocytes correlates with enhanced histopathological hallmarks of AD.

**Figure 2.**
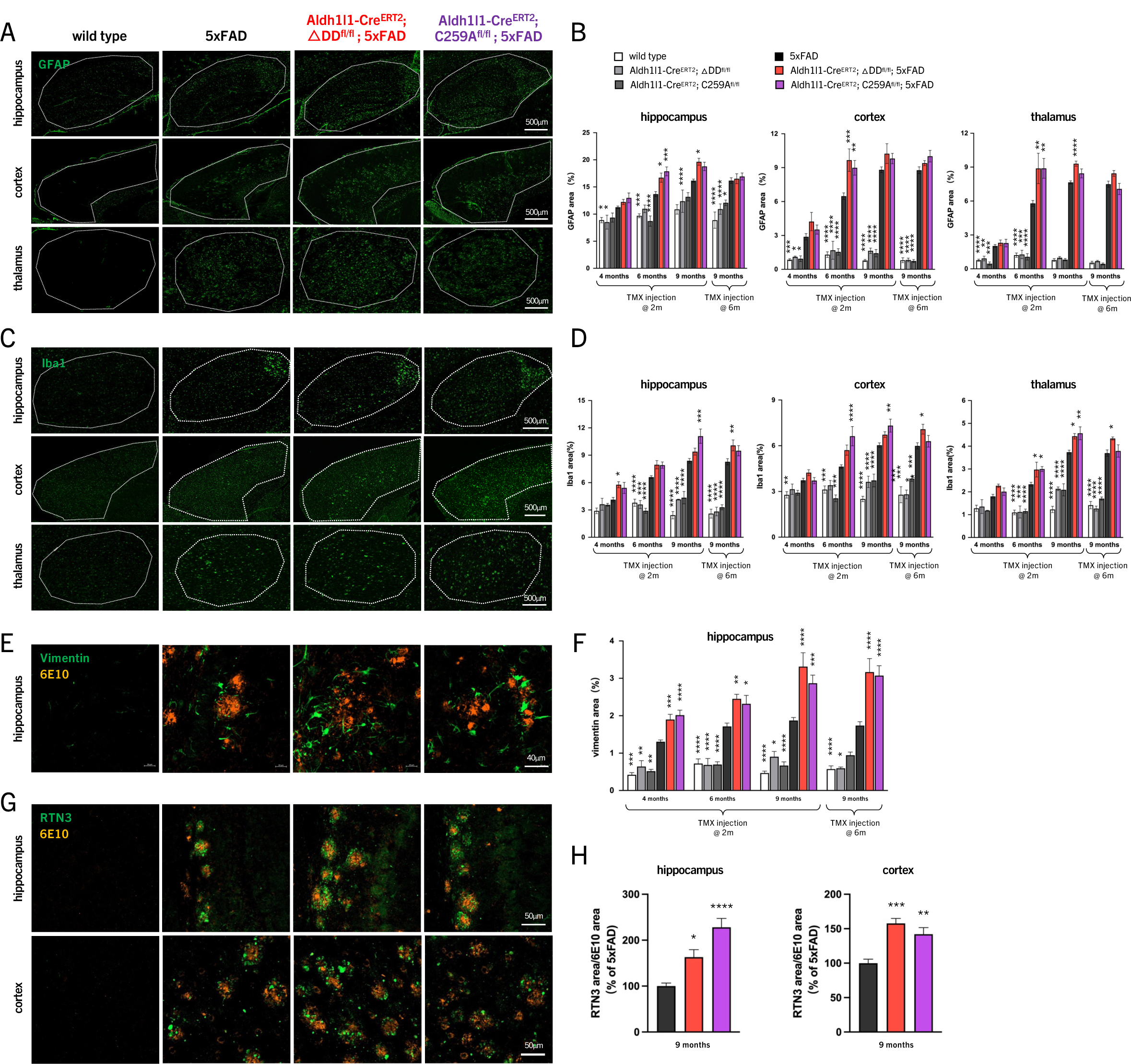
Increased AD brain histopathology in 5xFAD mice expressing signaling-deficient p75^NTR^ variants in astrocytes. (A) Representative micrographs showing GFAP immunostaining of sagittal sections through the hippocampus, cerebral cortex and thalamus of 5xFAD, Aldh1l1-CreER^T2^;ΔDD^fl/fl^;5xFAD and Aldh1l1-CreER^T2^;C259A^fl/fl^;5xFAD mice at 6 month of age. Mice were injected with tamoxifen at 2 month of age. The areas used for quantification are outlined. (B) Quantification of GFAP area (as % of outlined region) in hippocampus, cerebral cortex and thalamus of wild type, Aldh1l1-CreER^T2^;ΔDD^fl/fl^, Aldh1l1-CreER^T2^;C259A^fl/fl^, 5xFAD, Aldh1l1-CreER^T2^;ΔDD^fl/fl^;5xFAD and Aldh1l1-CreER^T2^;C259A^fl/fl^;5xFAD mice injected with TMX at 2 month or 6 month of age and analyzed at 4, 6 or 9 month as indicated. Results are presented as mean ± SEM (N=7-11 mice per group) and analyzed by two-way ANOVA followed by Tukey’s multiple comparisons test. *, p<0.05; **, p<0.01; ***, p<0.001; ****, p<0.0001 vs. 5xFAD of the corresponding age and condition. Histogram legend applies to all panels in this Figure. (C) Representative micrographs showing Iba1 immunostaining of sagittal sections through the hippocampus, cerebral cortex and thalamus of 5xFAD, Aldh1l1-CreER^T2^;ΔDD^fl/fl^;5xFAD and Aldh1l1-CreER^T2^;C259A^fl/fl^;5xFAD mice at 6 month of age. Mice were injected with tamoxifen at 2 month of age. The areas used for quantification are outlined. (D) Quantification of Iba1 area (as % of outlined region) in hippocampus, cerebral cortex and thalamus of wild type, Aldh1l1-CreER^T2^;ΔDD^fl/fl^, Aldh1l1-CreER^T2^;C259A^fl/fl^, 5xFAD, Aldh1l1-CreER^T2^;ΔDD^fl/fl^;5xFAD and Aldh1l1-CreER^T2^;C259A^fl/fl^;5xFAD mice injected with TMX at 2 month or 6 month of age and analyzed at 4, 6 or 9 month as indicated. Results are presented as mean ± SEM (N=7-11 mice per group) and analyzed by two-way ANOVA followed by Tukey’s multiple comparisons test. *, p<0.05; **, p<0.01; ***, p<0.001; ****, p<0.0001 vs. 5xFAD of the corresponding age and condition. (E) Representative micrographs showing Vimentin immunostaining of sagittal sections through the hippocampus of 5xFAD, Aldh1l1-CreER^T2^;ΔDD^fl/fl^;5xFAD and Aldh1l1-CreER^T2^;C259A^fl/fl^;5xFAD mice at 6 month of age. Mice were injected with tamoxifen at 2 month of age. (F) Quantification of Vimentin area (as % of region outlined in A) in hippocampus of wild type, Aldh1l1-CreER^T2^;ΔDD^fl/fl^, Aldh1l1-CreER^T2^;C259A^fl/fl^, 5xFAD, Aldh1l1-CreER^T2^;ΔDD^fl/fl^;5xFAD and Aldh1l1-CreER^T2^;C259A^fl/fl^;5xFAD mice injected with TMX at 2 month or 6 month of age and analyzed at 4, 6 or 9 month as indicated. Results are presented as mean ± SEM (N=7-11 mice per group) and analyzed by one-way ANOVA followed by Tukey’s multiple comparisons test. *, p<0.05; **, p<0.01; ***, p<0.001; ****, p<0.0001 vs. 5xFAD of the corresponding age and condition. (G) Representative micrographs showing RTN3 immunostaining of sagittal sections through the hippocampus and cerebral cortex of 5xFAD, Aldh1l1-CreER^T2^;ΔDD^fl/fl^;5xFAD and Aldh1l1-CreER^T2^;C259A^fl/fl^;5xFAD mice at 9 month of age. Mice were injected with tamoxifen at 2 month of age. (H) Quantification of RTN3 area (relative to Aβ plaque area) in hippocampus and cerebral cortex of 5xFAD, Aldh1l1-CreER^T2^;ΔDD^fl/fl^;5xFAD and Aldh1l1-CreER^T2^;C259A^fl/fl^;5xFAD mice injected with TMX at 2 month and analyzed at 9 month. Results (normalized to 5xFAD) are presented as mean ± SEM (N=7-11 mice per group) and analyzed by one-way ANOVA followed by Tukey’s multiple comparisons test. *, p<0.05; **, p<0.01; ***, p<0.001; ****, p<0.0001 vs. 5xFAD of the corresponding age and condition.

To assess whether the effects observed at the tissue level had an impact on cognitive performance, we compared wild type and 5xFAD mice bearing different p75^NTR^ variants in astrocytes. As 5xFAD mice are normally impaired in the classical Barnes maze test of hippocampal-dependent spatial learning and memory, we included brighter illumination and loud sound stimuli to increase the motivation of the mice to enter the escape chamber ^34,35^. This resulted in a partial learning and memory performance in these mice which, although still limited and below wild type levels, allowed us to test whether expression of signaling-deficient p75^NTR^ variants in astrocytes further affected these behaviors. Indeed, we found a significant worsening of learning performance in 5xFAD mice expressing mutant p75^NTR^ variants in astrocytes by the fourth training session compared to 5xFAD mice expressing wild type p75^NTR^ (Figure 3A). In the probe trails after the last training session (with escape chamber removed), 5xFAD mice carrying p75^NTR^ variants performed at the chance rate of 25%, below the level of 5xFAD mice expressing wild type p75^NTR^ (Figure 3B). Thus, the enhanced Aβ burden and brain histopathology displayed by AD mice with deficient p75^NTR^ signaling in astrocytes translates into measurable cognitive deficits.

**Figure 3.**
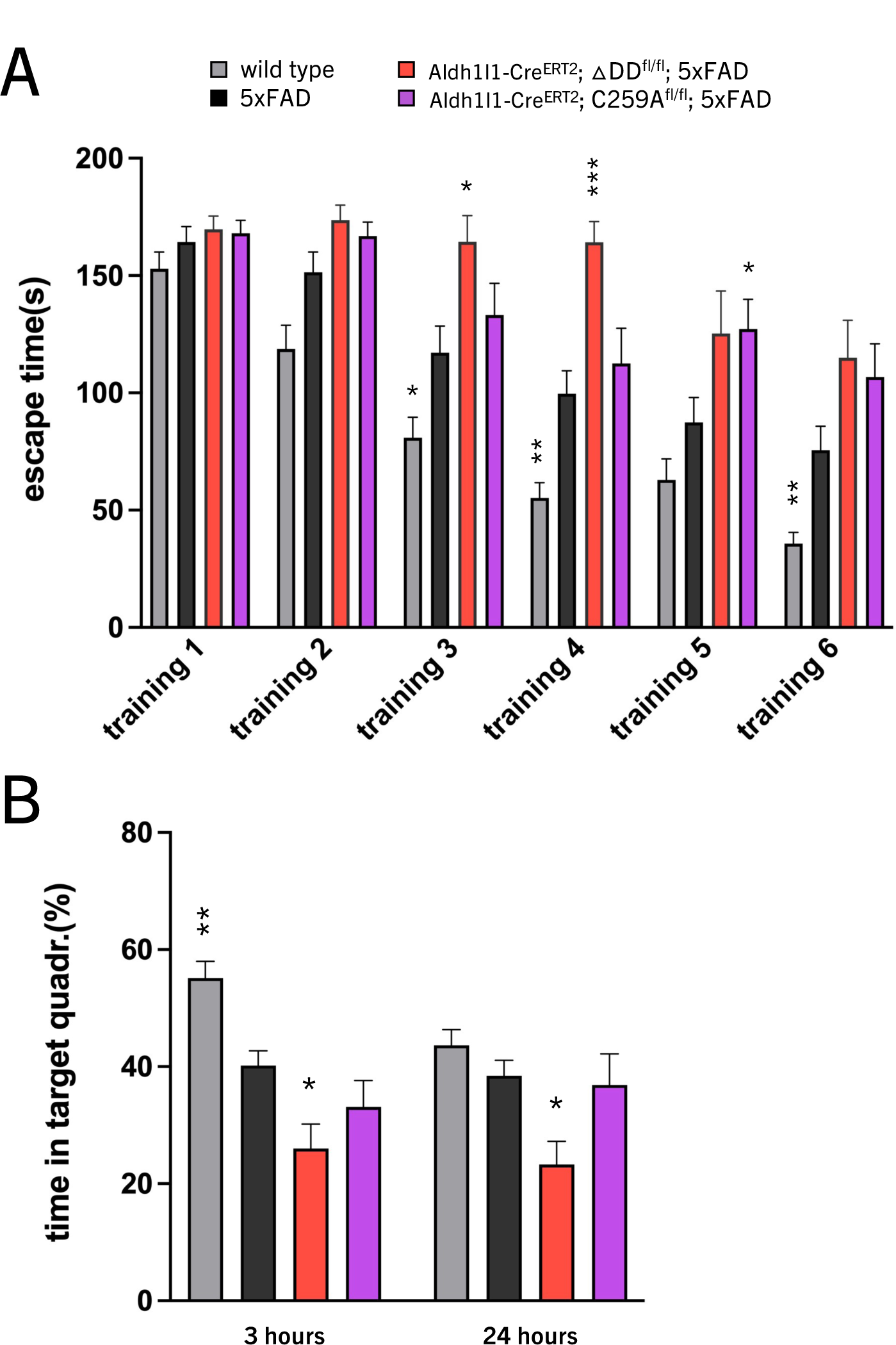
Increased learning/memory deficits in 5xFAD mice expressing signaling-deficient p75^NTR^ variants in astrocytes. (A) Training latency in the Barnes maze test of 6 month old wild type, 5xFAD, Aldh1l1-CreER^T2^;ΔDD^fl/fl^;5xFAD and Aldh1l1-CreER^T2^;C259A^fl/fl^;5xFAD mice. Results are shown as mean latency (in seconds) to find the platform hole ± SEM (N=16-20 mice per group) in 6 consecutive training sessions and analyzed by two -way ANOVA followed by Tukey’s multiple comparisons test. Bar color codes are as in panel (B). *, p<0.05; **, p<0.01; ***, p<0.001; ****, p<0.0001 vs. 5xFAD of the corresponding training day. Histogram legend applies to all panels in this Figure. (B) Percentage of time spent (mean ± SEM) in the target quadrant of the Barnes maze 3h and 24h after the last training session. Results are presented as mean ± SEM and analyzed by two-way ANOVA followed by Tukey’s multiple comparisons test. *, p<0.05; **, p<0.01 vs. 5xFAD of the corresponding test day. N numbers as in (A).

### Mutant astrocytes expressing signaling-deficient p75^NTR^ variants affect Aβ metabolism by both cell-autonomous and non-cell autonomous mechanisms

In the AD brain, astrocytes are known to participate in the clearance of Aβ plaques as well as influence neuronal production of Aβ ^2,3,36^. We examined the ability of cultured astrocytes bearing different p75^NTR^ variants to uptake preformed Aβ oligomers added to the culture medium as a measure of their ability to clear Aβ plaques from their environment. For these cultures, we isolated astrocytes from mice carrying constitutive ΔDD and C259A p75^NTR^ variants described in our earlier study ^6^. Following incubation with Aβ oligomers, astrocyte monolayers were washed and stained for Aβ and glial fibrillary protein (GFAP), an astrocyte marker, to assess engulfed Aβ aggregates. A significant reduction of Aβ aggregates associated with cell monolayers was observed in hippocampal and cortical astrocytes expressing ΔDD and C259A p75^NTR^ variants compared to wild type astrocytes (Figure 4A), suggesting that astrocytes expressing signaling-deficient p75^NTR^ variants have a diminished capacity to uptake Aβ oligomers. In agreement with a role for p75^NTR^ signaling in astrocyte Aβ oligomer uptake, treatment of wild type cortical and hippocampal astrocytes with p75^NTR^ activating ligands, such as NGF and BDNF, enhanced Aβ oligomer uptake (Figure 4B). Next, we tested the ability of astrocytes to affect Aβ production in wild type neurons by exposing the latter to conditioned medium from cultured astrocytes bearing different p75^NTR^ variants. Aβ was detected by ELISA in the media of neurons infected with a lentivirus expressing human APP carrying the same three mutations present in 5xFAD mice (Figure 4C). Conditioned medium from wild type astrocytes enhanced Aβ production in both cortical and hippocampus neurons by approximately two-fold (Figure 4C). Interestingly, conditioned medium from p75^NTR^ mutant astrocytes increased Aβ levels even further (Figure 4C), suggesting stronger non-cell-autonomous effects of astrocytes expressing signaling-deficient p75^NTR^ variants on neuronal Aβ production.

**Figure 4.**
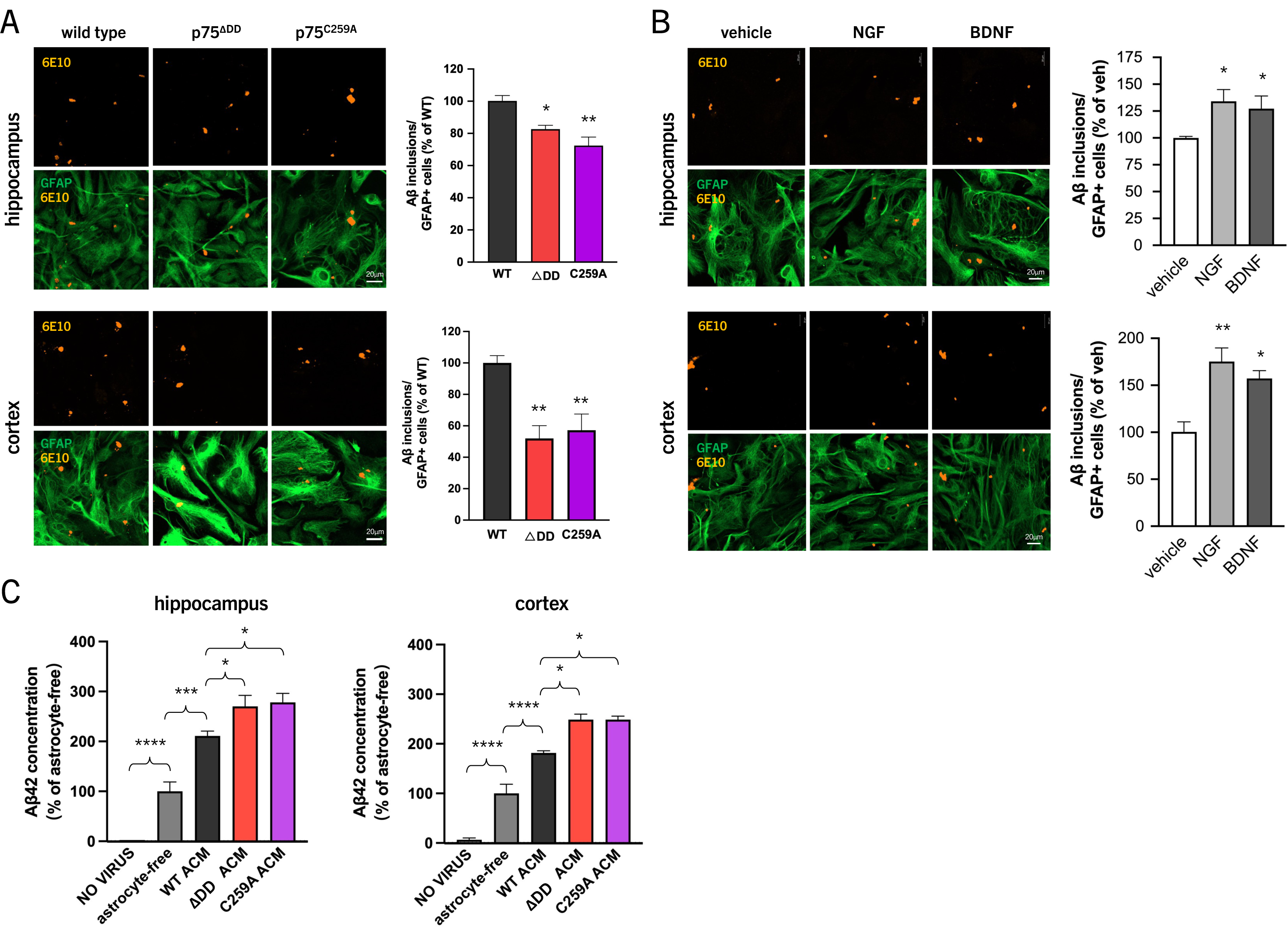
Mutant astrocytes expressing signaling-deficient p75^NTR^ variants affect Aβ metabolism by both cell-autonomous and non-cell autonomous mechanisms. (A) Representative micrographs of cultured astrocytes isolated from the hippocampus and cerebral cortex of wild type, p75^ΔDD^ and p75^C259A^ mice. Confluent monolayers were seeded with Aβ oligomers, washed and co-stained for Aβ (6E10) and GFAP to reveal uptake of Aβ plaques. Histograms show quantification of Aβ inclusions normalized to GFAP^+^ cell number and expressed as percentage of wild type (WT) levels. Results are presented as mean ± SEM (N=3 independent experiments each performed in triplicate) and analyzed by one-way ANOVA followed by Tukey’s multiple comparisons test. *, p<0.05; **, p<0.01 vs. WT. (B) Representative micrographs of cultured astrocytes isolated from the hippocampus and cerebral cortex of wild type mice. Confluent monolayers were treated with the indicated neurotrophins (50ng/ml), seeded with Aβ oligomers, washed and co-stained for Aβ (6E10) and GFAP to reveal uptake of Aβ plaques. Histograms show quantification of Aβ inclusions normalized to GFAP^+^ cell number and expressed as percentage of vehicle levels. Results are presented as mean ± SEM (N=3 independent experiments each performed in triplicate) and analyzed by one-way ANOVA followed by Tukey’s multiple comparisons test. *, p<0.05; **, p<0.01 vs. WT. (C) ELISA of Aβ-42 concentration in the supernatant of wild type hippocampal and cerebral cortex primary neuron cultures infected with AAV to express human AD mutant APP and treated with astrocyte conditioned medium (ACM) of the corresponding brain region derived from wild type, p75^ΔDD^ and p75^C259A^ mice as indicated. Results (mean ± SEM) are presented as percentage of astrocyte-free (N=3 independent experiments each performed in triplicate) and analyzed by one-way ANOVA followed by Tukey’s multiple comparisons test. *, p<0.05; ***, p<0.001; ****, p<0.0001 as indicated.

### Astrocyte expression of signaling-deficient p75^NTR^ variants up-regulates the cholesterol biosynthesis pathway

To investigate the molecular mechanisms by which astrocytes expressing signaling-deficient p75^NTR^ variants affect Aβ metabolism, we performed RNA-Seq studies in cultured hippocampal and cortical astrocytes isolated from new born mice constitutively expressing wild type or ΔDD or C259A p75^NTR^ variants ^6^. Candidate pathways selectively affected by the mutations were identified among gene sets (from Reactome and KEGG pathway databases) that were specifically upregulated in mutant compared to wild type astrocytes (Supplementary Figure S4). Among the top gene sets enriched in p75^NTR^ mutant astrocytes, “Cholesterol biosynthesis” and “Steroid biosynthesis” stood out as the only ones shared by ΔDD and C259A astrocytes from both brain regions (Supplementary Figure S4). These two gene sets contain most of the critical genes in the cholesterol biosynthesis pathway, including *Hmgcr* (encoding the rate-limiting enzyme of the mevalonate pathway), several *Dhcr* genes (encoding dehydrocholesterol reductases), *Fdft1* (encoding the enzyme catalyzing the first committed step in cholesterol biosynthesis), *Msmo1* (encoding a demethylase critical in cholesterol biosynthesis) and several others (Figure 5A). In line with this, increased protein levels of Sterol Regulatory Element-Binding Protein-2 (SREBP2), a master regulator of genes involved in cholesterol biosynthesis, and HMGCR were detected in cultured hippocampal and cortical ΔDD and C259A astrocytes compared to wild type (Figure 5B, C). Higher levels of SREBP2 were also found in hippocampal tissue extracts of 5xFAD mice expressing ΔDD and C259A p75^NTR^ variants in astrocytes compared to control or 5xFAD mice expressing wild type p75^NTR^ (Figure 5D). Together these results suggested the possibility that alterations in cholesterol biosynthesis underlay the enhanced AD neuropathology displayed by 5xFAD mice expressing signaling-deficient p75^NTR^ variants in astrocytes.

**Figure 5.**
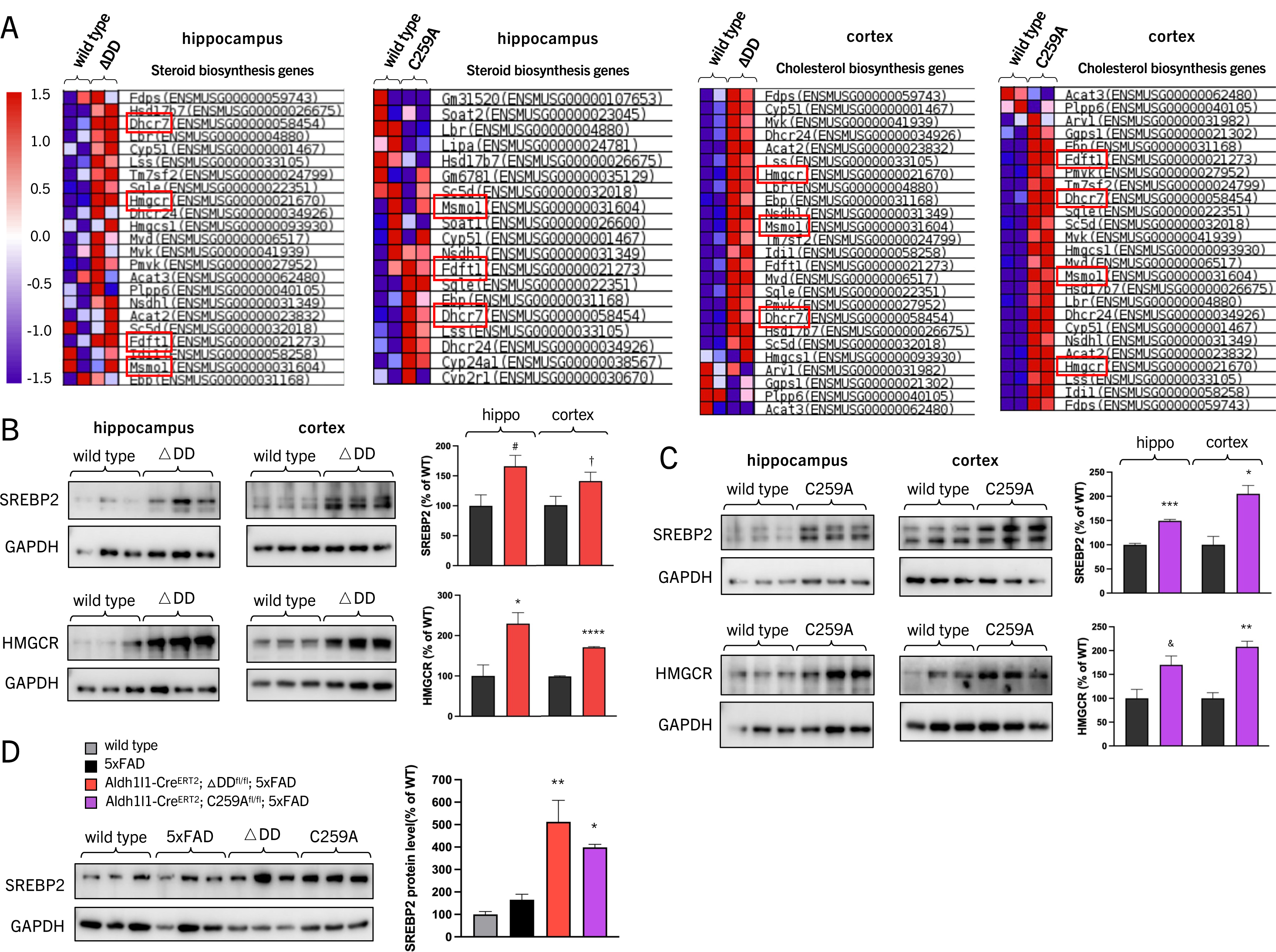
Astrocyte expression of signaling-deficient p75^NTR^ variants up-regulates the cholesterol biosynthesis pathway. (A) Genes in the steroid and cholesterol biosynthesis gene sets enriched in p75^ΔDD^ and p75^C259A^ hippocampal and cortical astrocytes as revealed by RNA-Seq. Results form two duplicate samples of each condition are shown. Selected genes *Dhcr7*, *Hmgcr*, *Fdft1* and *Msmo1* are highlighted. (B) Western blot analysis of SREBP2 and HMGCR expression in cultured hippocampal and cortical astrocytes from wild type and p75^ΔDD^ mice. Histograms show quantification of protein levels normalized to GAPDH and expressed as percentage of wild type (WT) levels. Results are presented as mean ± SEM (N=3) and analyzed by one-way ANOVA followed by Tukey’s multiple comparisons test. ^#^, p=0.062; †, p=0.12;*, p<0.05; **** p<0.0001 vs. WT. (C) Western blot analysis of SREBP2 and HMGCR expression in cultured hippocampal and cortical astrocytes from wild type and p75^C259A^ mice. Histograms show quantification of protein levels normalized to GAPDH and expressed as percentage of wild type (WT) levels. Results are presented as mean ± SEM (N=3 independent experiments each performed in triplicate) and analyzed by one-way ANOVA followed by Tukey’s multiple comparisons test. *, p<0.05; ***, p<0.001 vs. WT. (D) Western blot analysis of SREBP2 expression in hippocampal extracts from wild type, 5xFAD, Aldh1l1-CreER^T2^;ΔDD^fl/fl^;5xFAD and Aldh1l1-CreER^T2^;C259A^fl/fl^;5xFAD mice. Histogram show quantification of protein levels normalized to GAPDH and expressed as percentage of wild type (WT) levels. Results are presented as mean ± SEM (N=3 mice per group) and analyzed by one-way ANOVA followed by Tukey’s multiple comparisons test. *, p<0.05; **, p<0.01 vs. 5xFAD.

### Astrocyte expression of signaling-deficient p75^NTR^ variants increases cholesterol content in hippocampal astrocytes and neurons

In the adult brain, astrocytes are the main cell type capable of producing cholesterol, which they can transfer to neurons through Apolipoprotein E (ApoE) ^37–39^. Cholesterol has been shown to affect membrane fluidity and cell phagocytosis ^40–42^, as well as APP cleavage and Aβ production ^43,44^, and thus represented an excellent candidate for the mechanism underlying the cell-autonomous and non-cell autonomous effects of astrocytes on Aβ metabolism. In order to assess cholesterol levels in brain tissue and cultured cells, we employed filipin, a fluorescent, cholesterol binding antibiotic commonly used to detect and visualize free cholesterol. Elevated levels of cholesterol could be detected in hippocampal and cortical sections of mutant mice expressing signaling-deficient p75^NTR^ variants in astrocytes (Figure 6A and Supplementary Figure S5A). In agreement with a cell-autonomous upregulation of cholesterol levels, monolayers of ΔDD and C259A astrocytes showed increased filipin staining compared to wild type astrocytes (Figure 6B, and Supplementary Figure S5B). Conversely, cholesterol content in wild type hippocampal and cortical astrocytes was reduced upon treatment with p75^NTR^ activating ligands, such as NGF, proNGF and BDNF (Figure 6C and Supplementary Figure S5C). These ligands had no effect on the cholesterol content of ΔDD and C259A astrocytes (Supplementary Figure S6). Increased cholesterol levels were also detected in the conditioned medium of mutant astrocytes (Figure 6D and Supplementary Figure S5D); and wild type hippocampal and cortical neurons showed increased filipin staining after co-culture with mutant astrocytes compared to wild type astrocytes (Figure 6E and Supplementary Figure S5E).

**Figure 6.**
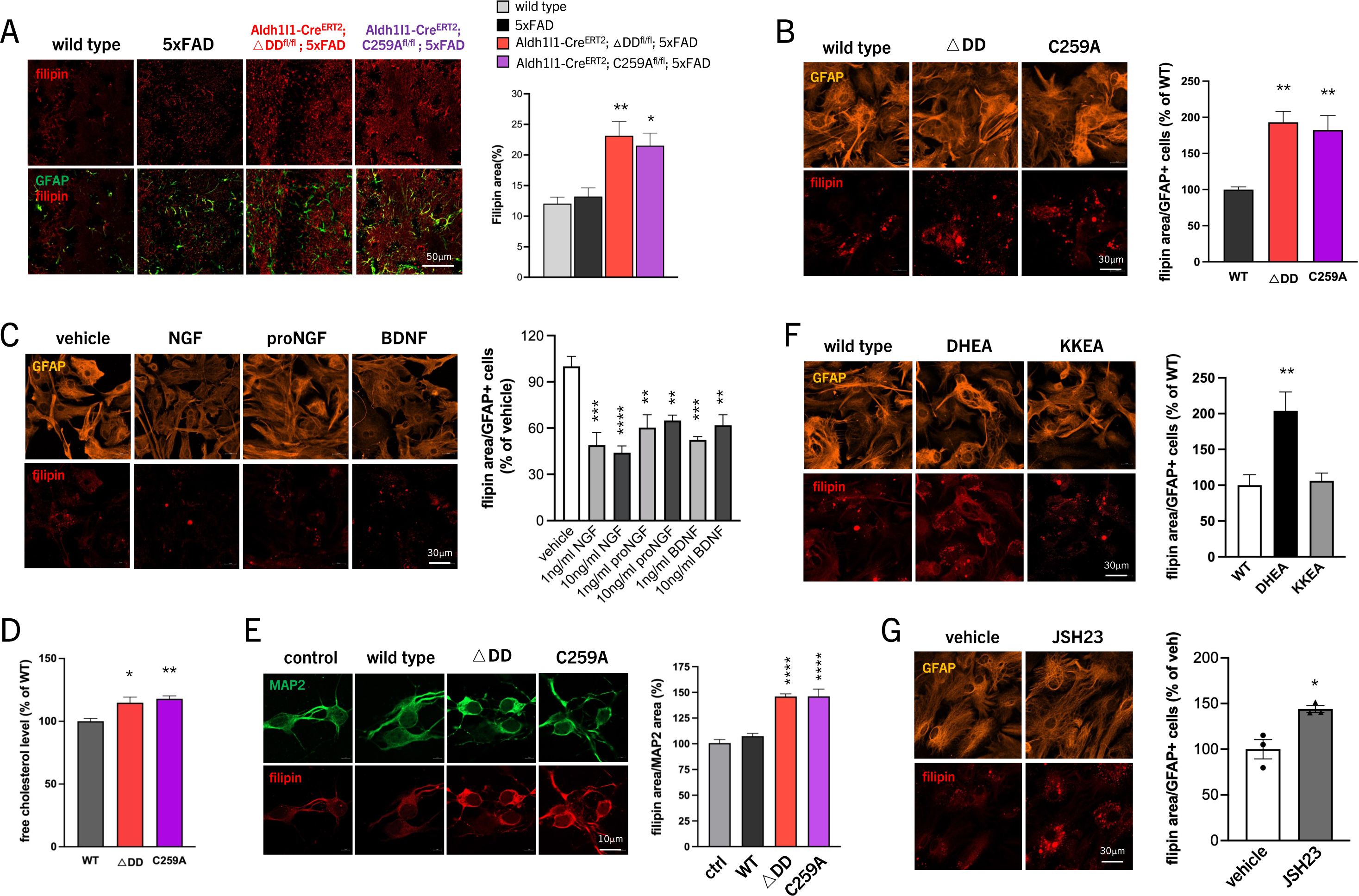
Astrocyte expression of signaling-deficient p75^NTR^ variants increases cholesterol content in hippocampal astrocytes and neurons. (A) Representative micrographs showing Filipin staining and GFAP immunostaining of sagittal sections through the hippocampus of wild type, 5xFAD, Aldh1l1-CreER^T2^;ΔDD^fl/fl^;5xFAD and Aldh1l1-CreER^T2^;C259A^fl/fl^;5xFAD mice at 6 month of age. Mice were injected with tamoxifen at 2 month of age. Histogram shows quantification of Filipin area (as % of GFAP area) in hippocampus of wild type, 5xFAD, Aldh1l1-CreER^T2^;ΔDD^fl/fl^;5xFAD and Aldh1l1-CreER^T2^;C259A^fl/fl^;5xFAD mice injected with TMX at 2 month and analyzed at 6 month. Results are presented as mean ± SEM (N=7-11 mice per group) and analyzed by one-way ANOVA followed by Tukey’s multiple comparisons test. *, p<0.05; **, p<0.01 vs. 5xFAD. (B) Representative micrographs of cultured astrocytes isolated from the hippocampus of wild type, p75^ΔDD^ and p75^C259A^ mice stained with Filipin to reveal cholesterol content and anti-GFAP antibody. Histogram shows quantification of Filipin area normalized to GFAP^+^ cell number and expressed as percentage of wild type (WT) levels. Results are presented as mean ± SEM (N=3 independent experiments each performed in triplicate) and analyzed by one-way ANOVA followed by Tukey’s multiple comparisons test. **, p<0.01 vs. WT. (C) Representative micrographs of cultured wild type hippocampal astrocytes treated with different neurotrophins at the indicated concentrations stained with Filipin to reveal cholesterol content and anti-GFAP antibody. Histogram shows quantification of Filipin area normalized to GFAP^+^ cell number and expressed as percentage of vehicle levels. Results are presented as mean ± SEM (N=3 independent experiments each performed in triplicate) and analyzed by one-way ANOVA followed by Tukey’s multiple comparisons test. **, p<0.01; ***, p<0.001; ****, p<0.0001 vs. vehicle. (D) Quantification of free cholesterol in the supernatant of cultured hippocampal astrocytes isolated from wild type, p75^ΔDD^ and p75^C259A^ mice. Results are presented as mean ± SEM (N=3 independent experiments each performed in triplicate) and analyzed by one-way ANOVA followed by Tukey’s multiple comparisons test. *, p<0.05; **, p<0.01 vs. WT. (E) Representative micrographs of cultured wild type hippocampal neurons supplemented with conditioned medium from wild type, p75^ΔDD^ and p75^C259A^ astrocytes stained with Filipin to reveal cholesterol content and anti-MAP2 antibody. Control indicates normal unsupplemented neuron culture medium. Histogram shows quantification of Filipin area normalized to MAP2 area and expressed as percentage of wild type (WT) levels. Results are presented as mean ± SEM (N=3 independent experiments each performed in triplicate) and analyzed by one-way ANOVA followed by Tukey’s multiple comparisons test. ****, p<0.0001 vs. control. (F) Representative micrographs of cultured astrocytes isolated from the hippocampus of wild type, p75^DHEA^ and p75^KKEA^ mice stained with Filipin to reveal cholesterol content and anti-GFAP antibody. Histogram shows quantification of Filipin area normalized to GFAP^+^ cell number and expressed as percentage of wild type (WT) levels. Results are presented as mean ± SEM (N=3 independent experiments each performed in triplicate) and analyzed by one-way ANOVA followed by Tukey’s multiple comparisons test. **, p<0.01 vs. WT. (G) Representative micrographs of cultured wild type hippocampal astrocytes treated with NF-kB inhibitor JSH23 (20ìM) stained with Filipin to reveal cholesterol content and anti-GFAP antibody. Histogram shows quantification of Filipin area normalized to GFAP^+^ cell number and expressed as percentage of vehicle levels. Results are presented as mean ± SEM (N=3 independent experiments each performed in triplicate) and analyzed by Student’s t-test. *, p<0.05 vs. vehicle.

These results suggested that p75^NTR^ signaling negatively regulates cholesterol biosynthesis in astrocytes, affecting the levels of cholesterol transferred to neurons. In order to gain an initial insight into the signaling pathways involved, we assessed cholesterol content in astrocytes derived from knock-in mutant mice carrying point mutations in the intracellular domain of p75^NTR^ that specifically interrupt coupling of the receptor to the NF-kB or RhoA pathways, two of the major signaling pathways downstream of p75^NTR^ (Supplementary Figure S7A, B). The triple replacement of Asp^357^, His^361^ and Glu^365^ with Ala in the death domain of p75^NTR^ (herein referred to as DHEA) prevents the binding of RIP2 ^45,46^, a critical adaptor protein linking the receptor to activation of NF-kB ^21^. On the other hand, Ala replacement of Lys^303^ in the juxtamembrane region together with Lys^346^ and Glu^349^ in the death domain (herein referred to as KKEA), abolishes the interaction of p75^NTR^ with RhoGDI and blunts its ability to regulate the RhoA pathway ^47^. We found that DHEA astrocytes, but not KKEA astrocytes, showed elevated cholesterol content as assessed by filipin staining (Figure 6F and Supplementary Figure S5F), mimicking the effects observed in ΔDD and C259A astrocytes. In line with this, treatment of wild type astrocytes with the NF-kB inhibitor JSH-23 increased filipin staining (Figure 6G and Supplementary Figure S5G). Together, these results suggested that p75^NTR^ signaling through the NF-kB pathway negatively regulates cholesterol biosynthesis in astrocytes.

### Cholesterol removal reverts cell-autonomous and non-cell autonomous effects of astrocyte signaling-deficient p75^NTR^ variants on Aβ metabolism

By altering membrane fluidity and compaction, increased cholesterol levels in mutant astrocytes could affect the ability of astrocytes to uptake Aβ oligomers. To test this idea, we treated astrocyte cultures with the cholesterol-quenching drug methyl-β-cyclodextrin (MβCD). Filipin staining of hippocampal and cortical astrocytes expressing p75^NTR^ variants ΔDD and C259A was reduced by approximately half following treatment with MβCD (Figure 7A), indicating reduced cholesterol content in the treated cells. Concomitantly, the ability of mutant astrocytes to uptake Aβ oligomers was enhanced by MβCD treatment, reaching levels comparable to those observed in wild type astrocytes (Figure 7B), suggesting that elevated cholesterol content impairs the ability of p75^NTR^ mutant astrocytes to clear Aβ plaques. Moreover, MβCD reduced Aβ production in cortical and hippocampal neurons expressing human APP carrying AD mutations (Figure 7C), demonstrating the direct relationship between cholesterol and Aβ production in AD neurons.

**Figure 7.**
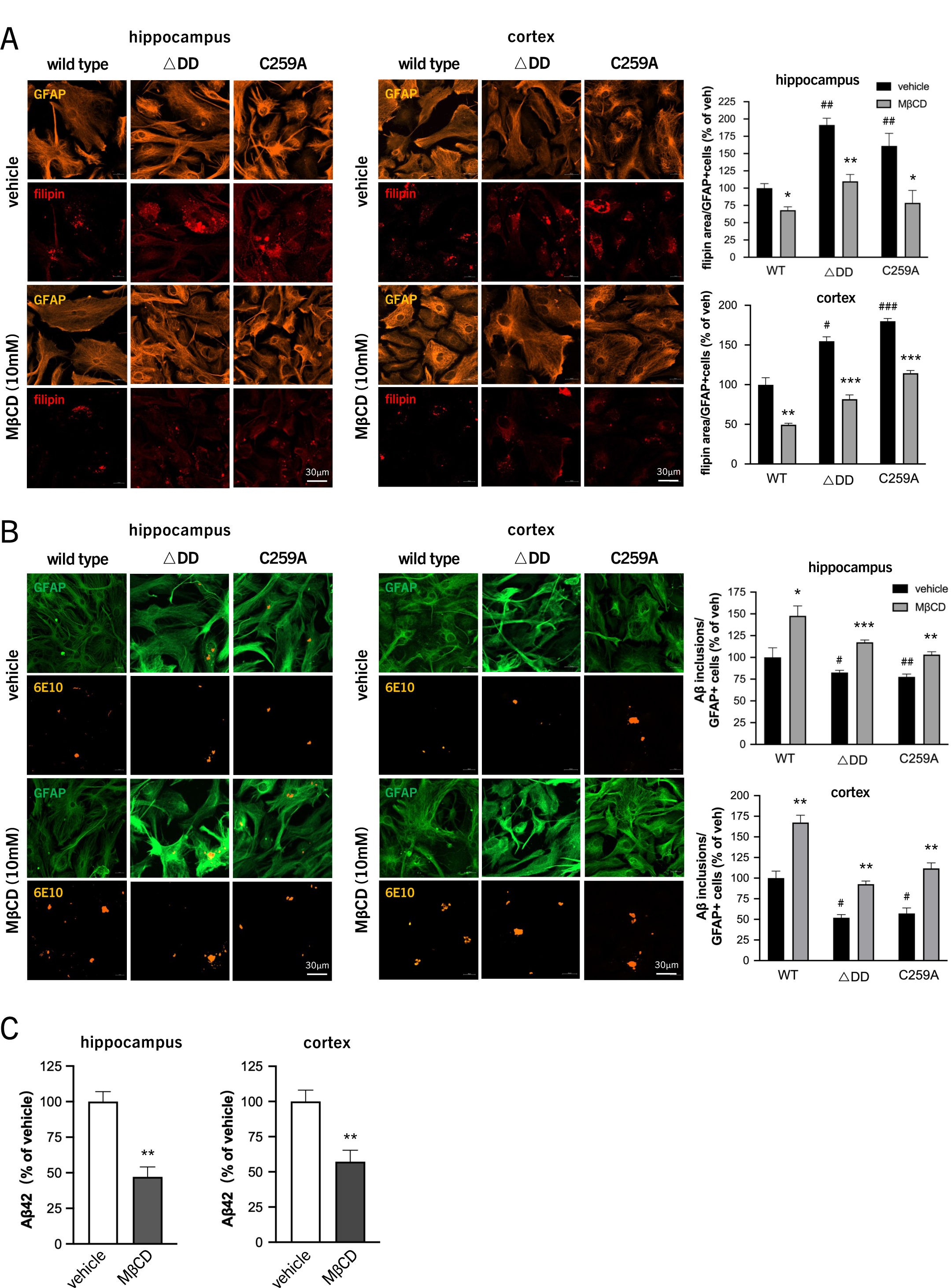
Cholesterol removal reverts cell-autonomous and non-cell autonomous effects of astrocyte signaling-deficient p75^NTR^ variants on Aβ metabolism. (A) Representative micrographs of cultured astrocytes isolated from the hippocampus and cerebral cortex of wild type, p75^ΔDD^ and p75^C259A^ mice treated with MβCD (10mM) or vehicle and stained with Filipin to reveal cholesterol content and anti-GFAP antibody. Histogram shows quantification of Filipin area normalized to GFAP^+^ cell number and expressed as percentage of vehicle-treated wild type (WT) levels. Results are presented as mean ± SEM (N=3 independent experiments each performed in triplicate) and analyzed by Student’s t-test. *, p<0.05; **, p<0.01: ***, p<0.001 vs. vehicle of corresponding genotype; #, p<0.05; ##, p<0.01; ###, p<0.001 vs. vehicle treated WT. (B) Representative micrographs of cultured astrocytes isolated from the hippocampus and cerebral cortex of wild type mice. Confluent monolayers were treated with MβCD (10mM) or vehicle, seeded with Aβ oligomers, washed and co-stained for Aβ (6E10) and GFAP to reveal uptake of Aβ plaques. Histograms show quantification of Aβ inclusions normalized to GFAP^+^ cell number and expressed as percentage of wild type (WT) levels. Results are presented as mean ± SEM (N=3 independent experiments each performed in triplicate) and analyzed by Student’s t-test. *, p<0.05; **, p<0.01: ***, p<0.001 vs. vehicle of corresponding genotype; #, p<0.05; ##, p<0.01 vs. vehicle treated WT. (C) ELISA of Aβ-42 concentration in the supernatant of wild type hippocampal and cerebral cortex primary neuron cultures infected with AAV to express human AD mutant APP and treated with MβCD (10mM) or vehicle. Results (mean ± SEM) are presented as percentage of vehicle (N=3 independent experiments each performed in triplicate) and analyzed by Student’s t-test. *, p<0.05 vs. vehicle.

### Statin treatment decreases brain cholesterol and counteracts the effects of impaired astrocyte p75^NTR^ function on AD neuropathology

Statins are powerful cholesterol lowering agents and broadly used in the treatment of arteriosclerosis and cardiovascular disease. In order to test the notion that signaling-deficient p75^NTR^ variants expressed in astrocytes enhance AD neuropathology by increasing brain cholesterol, we subjected 5xFAD mice expressing wild type or ΔDD p75^NTR^ in astrocytes to statin treatment. Following tamoxifen injection at 2 month of age, treatment with pravastatin administered through drinking water was initiated in 5xFAD (5xFAD::ΔDD^fl/fl^) and 5xFAD/ΔDD (5xFAD::*Aldh1l1-CreER*^T2^::ΔDD^fl/fl^) mutant mice. At 4 months of age, treatment with simvastatin administered by oral gavage was added and animals were sacrificed at 6 month (schematic in Figure 8A). Statin treatment significantly reduced cholesterol content in hippocampus and cortex of 5xFAD mice expressing ΔDD p75^NTR^ in astrocytes to levels comparable to those of 5xFAD mice expressing the wild type receptor (Figure 8B and Supplementary Figure S8A). Although statins also produced a small reduction in cholesterol levels in wild type and 5xFAD mice, those differences did not reach statistical significance. Analysis of Aβ burden revealed significantly reduced levels in hippocampus, cortex and thalamus of 5xFAD mice expressing ΔDD p75^NTR^ in astrocytes that were treated with statins (Figure 8C and Supplementary Figure S8B). A small reduction was also detected in 5xFAD expressing wild type p75^NTR^ but this only reached statistical significance in the thalamus (Figure 8C). Moreover, astrogliosis and microgliosis were also significantly decreased by statin treatment in all three brain regions of 5xFAD mice expressing ΔDD p75^NTR^ in astrocytes (Figure 8D, E and Supplementary Figure S8C, D). Again, statins also affected gliosis in 5xFAD mice expressing wild type p75^NTR^ but this did not reach statistical significance, while no effects were detected in control mice (Figure 8D, E). Thus, statin treatment effectively reverted the effects of astrocyte signaling-deficient p75^NTR^ variants on AD neuropathology, suggesting that they were primarily caused by increased brain cholesterol content as a result of enhanced astrocyte cholesterol biosynthesis.

**Figure 8.**
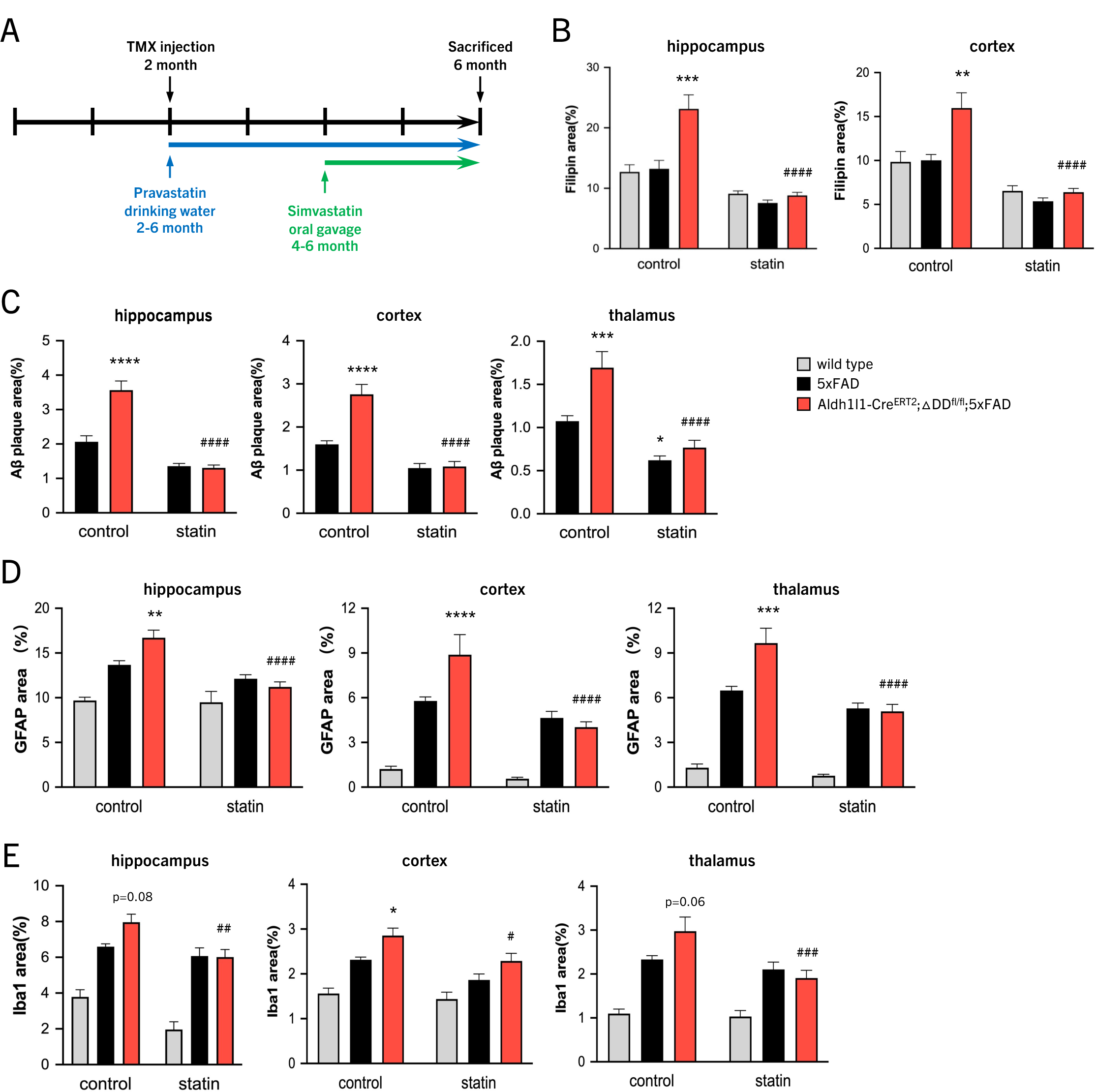
Statin treatment decreases brain cholesterol and counteracts the effects of impaired astrocyte p75^NTR^ function on AD neuropathology. (A) Schematic of timeline TMX injection and statin treatment. (B) Cholesterol content in hippocampus and cerebral cortex of wild type, 5xFAD and Aldh1l1-CreER^T2^;ΔDD^fl/fl^;5xFAD mice treated with vehicle (control) or statins assessed by Filipin staining (representative micrographs are shown in Figure S6). Histogram shows quantification of Filipin area (as % of GFAP area) presented as mean ± SEM (N=5-9 mice per group) and analyzed by two-way ANOVA followed by Tukey’s multiple comparisons test. **, p<0.01; ***, p<0.001 vs. vehicle-treated 5xFAD; ####, p<0.0001 vs. vehicle-treated Aldh1l1-CreER^T2^;ΔDD^fl/fl^;5xFAD. Histogram legend applies to all panels in this Figure. (C) Aâ plaque burden in hippocampus, cerebral cortex and thalamus of 5xFAD and Aldh1l1-CreER^T2^;ΔDD^fl/fl^;5xFAD mice treated with vehicle (control) or statins assessed by 6E10 immunostaining (representative micrographs are shown in Figure S6). Histogram shows quantification of Aβ area (as % of outlined region) presented as mean ± SEM (N=5-9 mice per group) and analyzed by two-way ANOVA followed by Tukey’s multiple comparisons test. *, p<0.05; ***, p<0.001 vs. vehicle-treated 5xFAD; ####, p<0.0001 vs. vehicle-treated Aldh1l1-CreER^T2^;ΔDD^fl/fl^;5xFAD. (D) Astrogliosis in hippocampus, cerebral cortex and thalamus of 5xFAD and Aldh1l1-CreER^T2^;ΔDD^fl/fl^;5xFAD mice treated with vehicle (control) or statins assessed by GFAP immunostaining (representative micrographs are shown in Figure S6). Histogram shows quantification of GFAP area (as % of outlined region) presented as mean ± SEM (N=5-9 mice per group) and analyzed by two-way ANOVA followed by Tukey’s multiple comparisons test. **, p<0.01; ***, p<0.001; ****, p<0.0001 vs. vehicle-treated 5xFAD; ####, p<0.0001 vs. vehicle-treated Aldh1l1-CreER^T2^;ΔDD^fl/fl^;5xFAD. (E) Microgliosis in hippocampus, cerebral cortex and thalamus of 5xFAD and Aldh1l1-CreER^T2^;ΔDD^fl/fl^;5xFAD mice treated with vehicle (control) or statins assessed by Iba1 immunostaining (representative micrographs are shown in Figure S8). Histogram shows quantification of Iba1 area (as % of outlined region) presented as mean ± SEM (N=5-9 mice per group) and analyzed by two-way ANOVA followed by Tukey’s multiple comparisons test. *, p<0.05 vs. vehicle-treated 5xFAD; ##, p<0.01; ###, p<0.001 vs. vehicle-treated Aldh1l1-CreER^T2^;ΔDD^fl/fl^;5xFAD.

## Discussion

All research studies performed to date on the role of p75^NTR^ in AD unanimously coincide on the capacity of this receptor to amplify the progression and pathological outcome of the disease. Impairment of p75^NTR^ function or expression, partial or complete, have invariably been reported to result in less severe pathology in mouse models of AD. Focusing mainly on neuronal p75^NTR^, these studies have attributed the detrimental effects of p75^NTR^ on AD pathology to enhanced receptor expression and coupling to apoptotic and neurodegenerative pathways, as well as increased APP intracellular trafficking and Aβ production. Although our present results do not challenge any of those findings, they do uncover a previously unknown function of p75^NTR^ in astrocytes that leads to a different outcome in AD, namely neuroprotection. We propose that such an effect results from the capacity of astrocyte p75^NTR^ to cell-autonomously dampen cholesterol biosynthesis and cholesterol secretion, thereby enhancing astrocyte Aβ uptake and reducing neuronal Aβ production. Our results also show that astrocyte p75^NTR^ counteracts AD pathology throughout the disease process, both at its onset and once established, indicating a continuous involvement in AD progression.

### Whole body vs. astrocyte-specific effects of signaling-deficient p75^NTR^ variants

The ΔDD and C259A mutations impair the ability of p75^NTR^ to properly respond to neurotrophins^29,30^. When introduced in a constitutive fashion from birth and in all cells, both result in significant neuroprotection in the 5xFAD mouse model of AD, intriguingly, at levels greater than blunting receptor expression altogether ^6^. By preserving receptor expression, and unlike knock-out models, signaling-deficient p75^NTR^ variants more closely parallel drug interventions that modify receptor function. This suggests that pharmacological strategies aiming at inhibiting receptor function, but preserving its expression, may be more beneficial than preventing receptor expression (as in siRNA-based therapies). Although p75^NTR^ is expressed in a variety of different cell types in the brain, including neurons, glial and endothelial cells, it has been assumed that its predominant function in AD was mainly restricted to neurons, as these are thought to be the main or sole cell type expressing APP in the brain. Several studies have shown such function to be deleterious, amplifying disease progression ^6–8^. Paradoxically, our present results indicate that, if the ΔDD and C259A mutations are introduced only in astrocytes, they lead to a significant acceleration and worsening of the disease. Unlike our previous whole-body study ^6^, in the present study the mutations were introduced in astrocytes from earliest 2 months of age. However, we have also shown here that cultured astrocytes isolated at early postnatal stages from constitutive mutant mice already display abnormal cholesterol biosynthesis and secretion, as well as reduced uptake of Aβ oligomers. It would therefore appear that, in AD, the beneficial effects afforded by impaired p75^NTR^ function across all receptor-expressing cells in the brain overcome the detrimental effects of impaired p75^NTR^ function in astrocytes. This is good news for ongoing efforts to manipulate p75^NTR^ function systemically in AD patients ^9^. Beyond this, our results also suggest that targeting the receptor in brain cells other than astrocytes, e.g. neurons, may afford even greater benefits in AD.

### p75^NTR^ signaling in cholesterol biosynthesis

Activation of receptor tyrosine kinases (RTKs) generally promote cholesterol biosynthesis by enhancing expression of SREBP2 as well as HMGCR, the rate-limiting step in the biosynthesis of cholesterol ^48–50^. Several studies have reported that p75^NTR^ can collaborate with RTKs to upregulate key steps in the cholesterol biosynthesis pathway. Thus, p75^NTR^ overexpression in melanoma cells was found to enhance the ability of TrkA to upregulate cholesterol biosynthesis ^51^. Similarly, NGF induces an increase in the expression of SREBP2 and HMGCR through p75^NTR^ and TrkA in PC12 cells ^52^; and p75^NTR^ can enhance the ability of the ErbB2 RTK to stimulate SREBP2 expression in Schwann cells ^53^. However, wherther p75^NTR^ is able to regulate cholesterol biosynthesis independently of RTK signaling has been less clear, particularly *in vivo* and in primary cells. Several lines of evidence suggest that the effects of astrocyte p75^NTR^ on cholesterol biosynthesis observed in the present study are likely mediated by p75^NTR^ independently of RTK signaling. First, astrocytes do not express full length Trk receptors bearing a tyrone kinase domain. Second, proNGF, a specific p75^NTR^ ligand that does not activate TrkA, reduced cholesterol content in astrocytes. Finally, neurotrophin ligands had no effects on the cholesterol content of ΔDD and C259A astrocytes. It should also be mentioned here that the effects of the ΔDD and C259A mutations on cholesterol biosynthesis and Aβ plaque uptake are unlikely to be the result of some gain-of-function in p75^NTR^ since all p75^NTR^ ligands tested showed opposite effects.

### NF-KB signaling in cholesterol biosynthesis

Among the best studied signaling cascades downstream of p75^NTR^ are the NF-kB and RhoA pathways. Using receptor mutants deficient in specific signaling pathways, we found that astrocytes expressing mutant p75^NTR^ deficient in coupling to the NF-kB pathway, but not those deficient in signaling to the RhoA pathway, showed increased cholesterol content similar to that observed in ΔDD and C259A astrocytes. This effect was mimicked by the NF-kB inhibitor JSH-23, and is in line with reports indicating that activation of NF-kB in macrophages by type I interferons or inflammatory substances such as lipopolysaccharide actively suppress cholesterol biosynthesis ^54–56^. It should be noted that NF-kB signaling has been shown to stimulate cholesterol content in hepatocytes and to upregulate SREBP2 expression in endothelial cells ^57,58^, suggesting that the effects of NF-kB signaling on cholesterol biosynthesis are cell-type specific.

### Cell-autonomous and non-cell-autonomous effects of astrocyte cholesterol in AD

Increased brain cholesterol levels can contribute to AD pathophysiology through different mechanisms (reviewed in ^44^. It has been shown that lipid membranes containing cholesterol can promote Aβ aggregation by enhancing its primary nucleation rate ^59^. In agreement with our findings, an early study found that cholesterol depletion attenuated Aβ generation in cultured hippocampal neurons ^60^. More recently, Barrett et al. reported that the transmembrane domain of APP can bind cholesterol, providing mechanistic insight into how cholesterol promotes amyloidogenesis ^61^. Related to this observation, is the finding that cholesterol can induce trafficking of APP to membrane compartments where it interacts with beta- and gamma-secretases to generate Aβ ^62^.

Beyond the effects of cholesterol in neurons, our study provides evidence on the cell-autonomous role of astrocyte p75^NTR^ in the regulation of cholesterol biosynthesis and the transfer of cholesterol to neurons, providing a key step in the cascade controlling brain cholesterol levels and its effects on Aβ production and aggregation. The inverse relationship between cholesterol content and the ability of astrocytes to uptake Aβ oligomers revealed by our results adds a previously unknown mechanism by which this lipid can negatively affect AD progression. The effects of cholesterol on membrane fluidity are bidirectional and likely complex ^40,41^. By altering the rigidity of the plasma membrane, cholesterol can affect the ability and rate at which cells phagocyte extracellular cargo^42^. Increased cholesterol content may also impair astrocyte uptake of Aβ aggregates by other mechanisms, including altered intracellular signaling or cytoskeletal plasticity.

### Statins, p75^NTR^ and AD therapeutics

Statins lower cholesterol levels by inhibiting HMGCR and are widely used in the treatment of cardiovascular disease. Despite abundant evidence from animal models ^63–65^, their effects on AD patients have been debated ^44,63^. Although early cohort and case-control studies indicated positive effects of statin treatment, subsequent randomized controlled trials have failed to establish a convincing link between statin treatment and cognitive improvement (reviewed in ^44^). Interestingly, one trial reported positive effects of statins in AD carriers of the ApoE4 allele ^66^, suggesting that further stratification of patient subpopulations according to their lipid homeostasis status could bring a fundamental breakthrough in the therapeutic use of cholesterol lowering drugs for AD. In our study, we used statins to test the hypothesis that expression of signaling-deficient variants of p75^NTR^ in astrocytes augment AD pathology by increasing cholesterol biosynthesis. Statin treatment reversed the enhanced AD pathology observed in these mutants, indicating that it was caused by increased brain cholesterol levels. A recent Phase II clinical trial in AD patients of a small molecule presented as a “modulator” of p75^NTR^ activity resulted in a mild, though not statistically significant, improvement in cognitive functions ^9^. It is not known which p75^NTR^ signaling pathways, if any, are affected by this drug. Our results suggest that selective interference with p75^NTR^ signaling pathways other than NF-kB may hold promise in the treatment of AD. Interestingly, a number of studies have implicated the RhoA pathway in Aβ production and aggregation, neuroinflammation and synaptic damage in AD (reviewed in ^67,68^. Our observation that KKEA astrocytes, expressing mutant p75^NTR^ deficient in coupling to the RhoA pathway, did not show elevated level of cholesterol, suggests that pharmacological approaches selectively targeting the ability of p75^NTR^ to affect RhoA signaling may hold promise in the treatment of AD.

## Methods

### Mice

Animal care and experimental procedures were approved by Laboratory Animal Welfare and Ethics Committee of Chinese Institute for Brain Research (CIBR-IACUC-028). Mice were housed in a 12-h light–dark cycle and fed a standard chow diet. The mouse lines utilized in this study are as follows: 5xFAD ^69^ Aldh1l1-Cre^ERT2^ ^31^; p75^NTR^ knock-out ^70^; conditional knock-in p75^NTR^ *ΔDD^fl^* and *C259A^fl^*, generated by Cyagen Co., China; constitutive knock-in p75^NTR^ ΔDD and C259A ^30^; constitutive knock-in p75^NTR^-DHEA, generated by Taconic-Artemis, Germany; and constitutive knock-in p75^NTR^-KKEA, generated at the Chinese Institute for Brain Research.

### Antibodies

The primary antibodies used for immunofluorescence (IF) and immunoblotting (WB) are as follows:

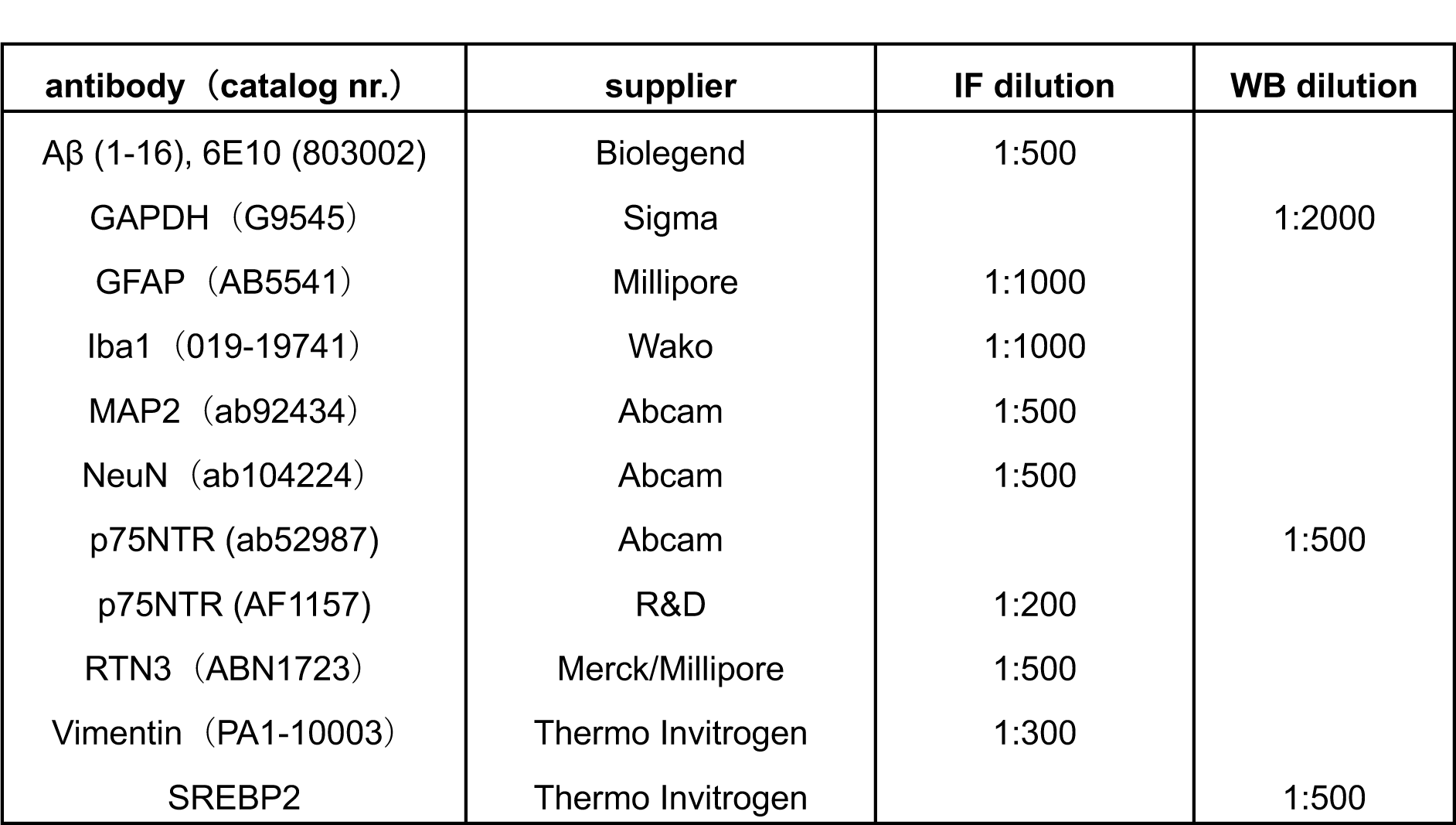

### Astrocyte culture and treatment

Astrocyte cultures were prepared from P2-P3 mouse pups. Cerebral cortex and hippocampal tissues were dissected aseptically from pups and enzymatically dissociated using 0.25% trypsin (Gibco) at 37°C for 25 minutes, which was terminated by complete growth medium. The digested tissues were mechanically triturated to generate single-cell suspensions. Cells were plated onto poly-D-lysine (Sigma)-coated T-75 flasks and maintained at 37°C in 5% CO₂ in Dulbecco’s Modified Eagle Medium (Gibco) supplied with 10% fetal bovine serum (BI) and 1% Penicillin/Streptomycin (Gibco). Upon reaching confluence after 7 days, microglia and oligodendrocyte were removed by shaking. Astrocytes were subsequently detached using 0.25% trypsin-EDTA (Gibco) for 5 minutes at 37°C, followed by terminated with complete growth medium. Cell density was counted using hemocytometer before replating.

Astrocytes were plated in the 24-well-plate at a density of 4 × 10^4^ cells/well onto PDL-coated coverslips until reaching confluency. For Aβ uptake assay, Aβ oligomers were prepared from Aβ1-42 peptide (Sigma), which were dissolved in PBS at a final concentration of 100 μM. The solution was vortexed vigorously for 1 min to ensure homogeneity and incubated at 37°C for 24 h. To remove insoluble aggregates or fibrils, the mixture was centrifuged at 14,000 × g for 10 min at 4°C. The supernatant containing soluble Aβ oligomers was collected, aliquoted kept in -20°C until use. Astrocytes were incubated with 0.5 uM Aβ oligomers for 24h at 37°C before washed with PBS and fixed. For neurotrophin treatment, different concentration of NGF (R&D), pro-NGF. (Alomone labs) and BDNF (Peprotech) were used and the cells were fixed 48h after the treatment. For MβCD treatment, astrocytes were pretreated with 2, 5 or 10 mM MβCD (Sigma) for 30 min before exposure to Aβ oligomers. For JSH-23 treatment, astrocytes were incubated with 20uM JSH-23 (Millipore) overnight before washed with PBS and fixed.

### Western blotting

Astrocyte and tissue lysates were prepared in TBST lysis buffer. The cell extract protein concentration was measured by a BCA Protein Assay Kit (Solarbio). Protein samples were mixed with 5X Laemmli buffer (Bio-rad), followed by boiled at 95°C for 5 minutes. 10% Tris-Tricine gels (Bio-Rad) were used. Proteins were then transferred to 0.2 μm polyvinylidene fluoride (PVDF) membranes (Amersham). Membranes were blocked with 3% non-fat milk milk (Bio-Rad) dissolved in TBST (Tris-buffered saline containing 0.1% Tween-20) for 1 hour at room temperature. Primary antibodies (Table 1) were applied in and incubated overnight at 4°C with gentle agitation. After three washes, membranes were probed with HRP-conjugated secondary antibodies diluted 1:2000 in 1% non-fat milk for 1 hour at room temperature. Immunoblots were developed using the chemiluminescent HRP substrate (Millipore) and exposed using ImageQuant (Amersham). Densitometric analysis was done using ImageJ software.

### Astrocyte bulk RNA sequencing and data analysis

Astrocytes were cultured until confluence as described above. Monolayers were lysed in TRIZOL and frozen at -80°C. Bulk RNA sequencing was outsourced to Novogene Co., Ltd, Beijing, China. Total amounts and integrity of RNA were assessed using the RNA Nano 6000 Assay Kit (Agilent Technologies) of the Bioanalyzer 2100 system. mRNA was enriched by oligo (dT) beads, fragmented using divalent cations under elevated temperature, and reverse-transcribed using random hexamer primer and M-MuLV Reverse Transcriptase, then use RNaseH to degrade the RNA. Second strand cDNA synthesis was performed using DNA Polymerase I and dNTP. After end repair and A-tailing, adaptor were ligated. Library fragments were purified with AMPure XP system (Beckman Coulter) to select cDNA fragments of 370∼420 bp. Then PCR amplification, the PCR product was purified by AMPure XP beads to obtain the library. Then the library was quantified by Qubit2.0 Fluorometer, and the insert size of the library is detected by Agilent 2100 bioanalyzer. qRT-PCR is used to accurately quantify the effective concentration of the library.

Sequencing was performed on Illumina NovaSeq 6000. Raw FASTQ data were processed through in-house perl. Scripts: removal of adapter, reads containing N base and low quality reads. Q20, Q30 and GC content the clean data were calculated. Then the clean reads were aligned to mm10 using Hisat2. The featureCounts v1.5.0-p3 was used to count the reads numbers mapped to each gene. Differential expression analysis was performed using the DESeq2 R package (1.20.0). Gene Ontology (GO), Reactome and KEGG pathways enrichment analysis of differentially expressed genes was implemented by the clusterProfiler R package (3.8.1). Gene Set Enrichment Analysis (GSEA) ranked the genes according to the degree of differential expression in the two samples, and then the predefined Gene Set were tested to see if they were enriched at the top or bottom of the list. Local version of the GSEA analysis tool http://www.broadinstitute.org/gsea/index.jsp was used to analyzed GO, KEGG data set.

### Neuron culture and treatment

Embryos were harvested from pregnant mice euthanized on gestational day 17 by cervical dislocation. Cerebral cortex and hippocampal tissues were dissected aseptically from embryos and enzymatically dissociated using Papain (Sigma-Aldrich) at 37°C for 25 minutes, which was terminated by horse serum. The digested tissues were mechanically triturated to generate single-cell suspensions and filtered through a 70μm strainer. Cell density was counted using hemocytometer before plating onto poly-D-lysine (Sigma) and Laminin (R&D)-coated coverslips. Cultures were maintained at 37°C in 5% CO₂ in Neurobasal medium (Gibco) supplemented with B27 (Thermo), GlutaMAX (Gibco), and 50 μg/ml Penicillin/Streptomycin (Gibco).

For neuron-astrocyte co-culture, astrocytes were cultured in 6-well plates at a density of 2×10⁵ cells/well in complete growth medium until reaching confluency. Cells were washed with PBS, and neuronal feeding medium was added 16h before co-culture. Neurons were plated in 24-well plates onto coated coverslips at 2×10⁵ cells/well in feeding medium. Neurons were pre-cultured for 4 days. For co-culture, neuron-loaded coverslips (neuronal side facing upward) were carefully transferred into the astrocyte-containing wells, ensuring no direct contact between the two cell types. After 2 days of co-culture, neuron-loaded coverslips were fixed with 4% PFA for 5 min at room temperature for filipin staining.

To obtain conditioned medium from astrocytes, astrocytes were plated in the 6-well-plate at a density of 2 × 10^5^ cells/well until reaching confluency. Cells were washed with PBS, and neuronal feeding medium was added. The medium was collected after 48 hours and centrifuged at 12, 000×g for 10 min to remove cellular debris. Neuronal cultures were treated with astrocyte-conditioned medium (ACM) from DIV3 to DIV9.

### AAV infection and Aβ ELISA

Neurons were treated with pAV-hSyn-hAPP virus (WZ biosciences Inc.) at DIV2 for 24h, which were then pretreated with 150 μM MβCD-cholesterol for 1 h or 5 mM MβCD for 30 min at DIV3. Then the cells were washed with PBS and fresh feeding media was added. To measure the effect of ACM on Aβ production, ACM was added at DIV3. The supernatant was collected at DIV9 for determination of the level of Aβ. The supernatant was collected and centrifuged at 12,000×g for 10 min to remove cell debris. Aβ1-42 levels were measured using a Human Amyloid 42 ELISA Kit (R&D) according to the manufacturer’s instructions.

### Immunocytochemistry, immunohistochemistry and filipin staining

Astrocytes or neurons cultured on coverslips were washed briefly in PBS, fixed with 4% PFA for 5 minutes at room temperature, and permeabilized and blocked in PBS containing 2% donkey serum and 0.2% Triton X-100 for 15min. After overnight incubation at 4°C with primary antibodies (see Table above), cells were washed and incubated with Alexa Fluor-conjugated donkey anti-rabbit/mouse/chicken IgG secondary antibodies for 2 hours at room temperature, followed by mounting with fluorescence mounting medium (Dako) and imaging on a LSM 900 confocal microscope (Zeiss).

Mice were anesthetized with isoflurane and perfused with PBS until the liver turned pale. Then the brain tissue was extracted and divided into two hemispheres. The left hemisphere was fixed in 4% PFA at 4°C for 16 hours, washed three times with PBS, and then sequentially dehydrated in 20% and 30% sucrose solutions at 4°C until it sank to the bottom, after which the left hemisphere was embedded in OCT, sagittally sectioned at a thickness of 14 µm using a cryostat. For the right hemisphere, the hippocampus and cortex were dissected out and stored at -80°C.

For immunohistochemistry, cryostat sections (14 μm) were permeabilized and blocked in PBS containing 2% donkey serum and 0.5% Triton X-100 for 15min. Sections were stained with primary antibodies (see Table above) overnight at 4°C, washed, and incubated with Alexa Fluor-conjugated donkey anti-rabbit/mouse/chicken IgG secondary antibodies for 2 hours at room temperature before mounting and imaging on a LSM 900 confocal microscope.

For filipin staining, neurons or astrocytes placed on the coverslips were fixed with 4% PFA for 5 min at room temperature. After permeabilization with 0.2% Triton X-100 for 15min, the cells were stained with 20 μg/mL filipin (Sigma) in PBS for 1 h at room temperature before mounting and imaging on a LSM 900 confocal microscope. Brain sections were permeabilization with 0.5% Triton X-100 for 15min, then stained with 40 μg/mL filipin in PBS for 1 h at room temperature before mounting and imaging on a LSM 900 confocal microscope.

### Cholesterol secretion assay

Astrocytes were plated in the 24-well-plate at a density of 4 × 104 cells/well until reaching confluency. The cells were kept for 3 days in serum-free medium, then the supernatant was collected and centrifuged at 12,000 g for 10 minutes to remove debris. The amount of cholesterol in the medium was measured using cholesterol quantitation Kit (Sigma). The working solution contains of 300 μM Amplex Red reagent, 2 U/mL HRP, 2 U/mL cholesterol oxidase, and 0.2 U/mL cholesterol esterase. In a 96-well plate, 50 µl of the supernatant and 50 µl of the working solution were added to each well and incubate for 15 min at 37°C. The concentration of cholesterol was determined by an enzyme-coupled reaction. Cholesterol was oxidized by cholesterol oxidase to yield H_2_O_2_, which was then detected using the Amplex Red reagent. In the presence of HRP, Amplex Red reagent reacts with H_2_O_2_ to produce highly fluorescent resorufin, which was then measured using excitation at 540 nm and emission detection at 590 nm in microplate reader (BioTek).

### Barnes maze

The Barnes maze was conducted using a circular platform with an escape tunnel underneath. Spatial cues were placed around the platform. On the first day of training, the mouse (6 month old) was first placed in the escape tunnel for 1 minute. The animal was then placed in the center of the maze inside a box, which was removed after 15 seconds as in all subsequent sessions. The mouse was then allowed to explore the maze for 3 minutes or until it found the escape tunnel. To avoid the odorant cue, the escape platform was rotated 90° daily while the spatial cues and tunnel location remained the same. Mice were trained once daily for 6 days. For the memory test, the escape tunnel was removed and mouse was free to explore the maze for 90 seconds. Short-term (2h post-training) memory and long-term (24h post-training) memory were measured by quantifying the time spent in the target quadrant. 80dB noise and 500lux light were applied on all trials.

### Statin treatment

Two month old male mice were used in these experiments. Pravastatin was added in the drinking water, which was administrated at 100mg/L for 3 d, increased to 150mg/L for 4 d, and then increased to 200mg/L for the rest of the treatment. Simvastatin was administrated intraperitoneally starting at 4 month of age, at a concentration of 40 mg/kg/d, three times a week. Mice were sacrificed at 6 month of age.

### Statistical analysis

Statistical Analysis Software (SAS Institute, Inc.) and GraphPad Prism (versions 4 or 8; GraphPad Software Inc, San Diego, CA, USA) were used for statistical analyses. Experimental data was collected from multiple experiments and reported as the treatment mean ± SEM. Statistical significance was calculated using the one-way ANOVA or two-way ANOVA followed by Tukey’s multiple comparisons test for all data except that shown in Figures 7C, S3B and S5G which used the Student’s t-test. P value of less than 0.05 was considered statistically significant.

## Acknowledgements

The authors would like to thank Lei Wang, Jocelyn Jia, Yankui Fu and Shuo Zhang for technical and admin assistance. This work was supported by research grants to C.F.I. from Peking University, Chinese Institute for Brain Research, Beijing, and Swedish Research Council (Vetenskapsrådet, contract nr. 2024-03222); and a startup grant to M.X. from Swedish Research Council (Vetenskapsrådet, contract nr. 2021-01805).

## Author contributions

X.H. performed all experimental work, analyzed data and prepared a draft of the figures; M.X. co-directed the project and corrected the manuscript; C.F.I. conceived the project, directed the research, and wrote the manuscript.

## References

1. Strooper, B. D. & Karran, E. The Cellular Phase of Alzheimer’s Disease. Cell 164, 603–615 (2016).

2. Carter, S. F. et al. Astrocyte Biomarkers in Alzheimer’s Disease. Trends Mol Med 25, 77–95 (2019).

3. Qin, H. et al. Diverse signaling mechanisms and heterogeneity of astrocyte reactivity in Alzheimer’s disease. J. Neurochem. (2023) doi:10.1111/jnc.16002.

4. Burda, J. E. & Sofroniew, M. V. Reactive Gliosis and the Multicellular Response to CNS Damage and Disease. Neuron 81, 229–248 (2014).

5. Fombonne, J., Rabizadeh, S., Banwait, S., Mehlen, P. & Bredesen, D. E. Selective vulnerability in Alzheimer’s disease: amyloid precursor protein and p75(NTR) interaction. Ann. Neurol. 65, 294– 303 (2009).

6. Yi, C. et al. Inactive variants of death receptor p75NTR reduce Alzheimer’s neuropathology by interfering with APP internalization. Embo J 40, e104450 (2021).

7. Wang, Y.-J. et al. p75NTR regulates Abeta deposition by increasing Abeta production but inhibiting Abeta aggregation with its extracellular domain. J Neurosci 31, 2292–2304 (2011).

8. Knowles, J. K. et al. The p75 Neurotrophin Receptor Promotes Amyloid-beta(1-42)-Induced Neuritic Dystrophy In Vitro and In Vivo. J Neurosci 29, 10627–10637 (2009).

9. Shanks, H. R. C. et al. p75 neurotrophin receptor modulation in mild to moderate Alzheimer disease: a randomized, placebo-controlled phase 2a trial. Nat. Med. 30, 1761–1770 (2024).

10. Ernfors, P., Lindefors, N., Chan-Palay, V. & Persson, H. Cholinergic neurons of the nucleus basalis express elevated levels of nerve growth factor receptor mRNA in senile dementia of the Alzheimer type. Dementia 1, 138–145 (1990).

11. Mufson, E. J. & Kordower, J. H. Cortical neurons express nerve growth factor receptors in advanced age and Alzheimer disease. Proc Natl Acad Sci USA 89, 569–573 (1992).

12. Hu, X.-Y. et al. Increased p75(NTR) expression in hippocampal neurons containing hyperphosphorylated tau in Alzheimer patients. Exp Neurol 178, 104–111 (2002).

13. Chakravarthy, B. et al. Hippocampal membrane-associated p75NTR levels are increased in Alzheimer’s disease. J. Alzheimers Dis. 30, 675–684 (2012).

14. Chakravarthy, B. et al. Amyloid-beta peptides stimulate the expression of the p75(NTR) neurotrophin receptor in SHSY5Y human neuroblastoma cells and AD transgenic mice. J. Alzheimers Dis. 19, 915–925 (2010).

15. Underwood, C. K. & Coulson, E. J. The p75 neurotrophin receptor. Int J Biochem Cell Biol 40, 1664–1668 (2008).

16. Ibáñez, C. F. & Simi, A. p75 neurotrophin receptor signaling in nervous system injury and degeneration: paradox and opportunity. Trends Neurosci 35, 431–440 (2012).

17. Friedman, W. J. Neurotrophins induce death of hippocampal neurons via the p75 receptor. J Neurosci 20, 6340–6346 (2000).

18. Carter, B. D. et al. Selective activation of NF-kappa B by nerve growth factor through the neurotrophin receptor p75. Science 272, 542–545 (1996).

19. Yamashita, T., Higuchi, H. & Tohyama, M. The p75 receptor transduces the signal from myelin-associated glycoprotein to Rho. J Cell Biol 157, 565–570 (2002).

20. Yamashita, T., Tucker, K. L. & Barde, Y.-A. Neurotrophin binding to the p75 receptor modulates Rho activity and axonal outgrowth. Neuron 24, 585–593 (1999).

21. Khursigara, G. et al. A prosurvival function for the p75 receptor death domain mediated via the caspase recruitment domain receptor-interacting protein 2. J. Neurosci. : Off. J. Soc. Neurosci. 21, 5854–63 (2001).

22. Patapoutian, A. & Reichardt, L. F. Trk receptors: mediators of neurotrophin action. Curr Opin Neurobiol 11, 272–280 (2001).

23. Bothwell, M. Recent advances in understanding context-dependent mechanisms controlling neurotrophin signaling and function. F1000Research 8, F1000 Faculty Rev-1658 (2019).

24. Volosin, M. et al. Interaction of survival and death signaling in basal forebrain neurons: roles of neurotrophins and proneurotrophins. J Neurosci 26, 7756–7766 (2006).

25. Cragnolini, A. B., Huang, Y., Gokina, P. & Friedman, W. J. Nerve growth factor attenuates proliferation of astrocytes via the p75 neurotrophin receptor. Glia 57, 1386–1392 (2009).

26. Choi, S. & Friedman, W. J. Inflammatory cytokines IL-1â and TNF-á regulate p75NTR expression in CNS neurons and astrocytes by distinct cell-type-specific signalling mechanisms. ASN Neuro 1, (2009).

27. Capsoni, S. et al. The chemokine CXCL12 mediates the anti-amyloidogenic action of painless human nerve growth factor. Brain 140, 201–217 (2017).

28. Domeniconi, M., Hempstead, B. L. & Chao, M. V. Pro-NGF secreted by astrocytes promotes motor neuron cell death. Mol Cell Neurosci 34, 271–279 (2007).

29. Vilar, M. et al. Activation of the p75 neurotrophin receptor through conformational rearrangement of disulphide-linked receptor dimers. Neuron 62, 72–83 (2009).

30. Tanaka, K., Kelly, C. E., Goh, K. Y., Lim, K. B. & Ibáñez, C. F. Death Domain Signaling by Disulfide-Linked Dimers of the p75 Neurotrophin Receptor Mediates Neuronal Death in the CNS. J Neurosci 36, 5587–5595 (2016).

31. Hu, N.-Y. et al. Expression Patterns of Inducible Cre Recombinase Driven by Differential Astrocyte-Specific Promoters in Transgenic Mouse Lines. Neurosci Bull 36, 530–544 (2020).

32. Habib, N. et al. Disease-associated astrocytes in Alzheimer’s disease and aging. Nat Neurosci 23, 701–706 (2020).

33. Hu, X. et al. Transgenic mice overexpressing reticulon 3 develop neuritic abnormalities. EMBO J 26, 2755–2767 (2007).

34. Pitts, M. Barnes Maze Procedure for Spatial Learning and Memory in Mice. BIO-Protoc. 8, (2018).

35. Ingersoll, J. et al. Analyzing Spatial Learning and Prosocial Behavior in Mice Using the Barnes Maze and Damsel-in-Distress Paradigms. J. Vis. Exp. (2018) doi:10.3791/58008.

36. Sarkar, S. & Biswas, S. C. Astrocyte subtype-specific approach to Alzheimer’s disease treatment. Neurochem. Int. 145, 104956 (2021).

37. Dietschy, J. M. & Turley, S. D. Cholesterol metabolism in the central nervous system during early development and in the mature animal. J. Lipid Res. 45, 1375–1397 (2004).

38. Russell, D. W., Halford, R. W., Ramirez, D. M. O., Shah, R. & Kotti, T. Cholesterol 24-Hydroxylase: An Enzyme of Cholesterol Turnover in the Brain. Annu. Rev. Biochem. 78, 1017– 1040 (2009).

39. Vance, J. E. & Hayashi, H. Formation and function of apolipoprotein E-containing lipoproteins in the nervous system. Biochim. Biophys. Acta (BBA) - Mol. Cell Biol. Lipids 1801, 806–818 (2010).

40. Zhang, Y., Li, Q., Dong, M. & Han, X. Effect of cholesterol on the fluidity of supported lipid bilayers. Colloids Surf. B: Biointerfaces 196, 111353 (2020).

41. Pöhnl, M., Trollmann, M. F. W. & Böckmann, R. A. Nonuniversal impact of cholesterol on membranes mobility, curvature sensing and elasticity. Nat. Commun. 14, 8038 (2023).

42. Saharan, O., Mehendale, N. & Kamat, S. S. Phagocytosis: A (Sphingo)Lipid Story. Curr. Res. Chem. Biol. 2, 100030 (2022).

43. Wolozin, B. Cholesterol and the Biology of Alzheimer’s Disease. Neuron 41, 7–10 (2004).

44. Ahmed, H. et al. Brain cholesterol and Alzheimer’s disease: challenges and opportunities in probe and drug development. Brain 147, 1622–1635 (2024).

45. Charalampopoulos, I. et al. Genetic dissection of neurotrophin signaling through the p75 neurotrophin receptor. Cell Rep 2, 1563–1570 (2012).

46. Vicario, A., Kisiswa, L., Tann, J. Y., Kelly, C. E. & Ibáñez, C. F. Neuron-type-specific signaling by the p75NTR death receptor is regulated by differential proteolytic cleavage. J Cell Sci 128, 1507–1517 (2015).

47. Ramanujan, A. & Ibáñez, C. F. RhoGDI phosphorylation by PKC promotes its interaction with death receptor p75 NTR to gate axon growth and neuron survival. EMBO Rep. 23-Jan (2024) doi:10.1038/s44319-024-00064-2.

48. Bengoechea-Alonso, M. T. & Ericsson, J. SREBP in signal transduction: cholesterol metabolism and beyond. Curr. Opin. Cell Biol. 19, 215–222 (2007).

49. Porstmann, T. et al. SREBP Activity Is Regulated by mTORC1 and Contributes to Akt-Dependent Cell Growth. Cell Metab. 8, 224–236 (2008).

50. Clendening, J. W. et al. Exploiting the mevalonate pathway to distinguish statin-sensitive multiple myeloma. Blood 115, 4787–4797 (2010).

51. Restivo, G. et al. The low neurotrophin receptor CD271 regulates phenotype switching in melanoma. Nat Commun 8, 1988 (2017).

52. Comaposada-Baró, R. et al. Cholinergic neurodegeneration and cholesterol metabolism dysregulation by constitutive p75NTR signaling in the p75exonIII-KO mice. Front. Mol. Neurosci. 16, 1237458 (2023).

53. Follis, R. M. et al. Metabolic Control of Sensory Neuron Survival by the p75 Neurotrophin Receptor in Schwann Cells. J Neurosci 41, 8710–8724 (2021).

54. York, A. G. et al. Limiting Cholesterol Biosynthetic Flux Spontaneously Engages Type I IFN Signaling. Cell 163, 1716–1729 (2015).

55. Dang, E. V., McDonald, J. G., Russell, D. W. & Cyster, J. G. Oxysterol Restraint of Cholesterol Synthesis Prevents AIM2 Inflammasome Activation. Cell 171, 1057–1071.e11 (2017).

56. Araldi, E. et al. Lanosterol Modulates TLR4-Mediated Innate Immune Responses in Macrophages. Cell Rep. 19, 2743–2755 (2017).

57. Fowler, J. W. M., Zhang, R., Tao, B., Boutagy, N. E. & Sessa, W. C. Inflammatory stress signaling via NF-kB alters accessible cholesterol to upregulate SREBP2 transcriptional activity in endothelial cells. eLife 11, e79529 (2022).

58. Heida, A. et al. The hepatocyte IKK:NF-κB axis promotes liver steatosis by stimulating de novo lipogenesis and cholesterol synthesis. Mol. Metab. 54, 101349 (2021).

59. Habchi, J. et al. Cholesterol catalyses Aβ42 aggregation through a heterogeneous nucleation pathway in the presence of lipid membranes. Nat. Chem. 10, 673–683 (2018).

60. Simons, M. et al. Cholesterol depletion inhibits the generation of β-amyloid in hippocampal neurons. Proc. Natl. Acad. Sci. 95, 6460–6464 (1998).

61. Barrett, P. J. et al. The amyloid precursor protein has a flexible transmembrane domain and binds cholesterol. Science 336, 1168–1171 (2012).

62. Wang, H. et al. Regulation of beta-amyloid production in neurons by astrocyte-derived cholesterol. P Natl Acad Sci Usa 118, e2102191118 (2021).

63. Rahman, S. O., Hussain, S., Alzahrani, A., Akhtar, Mohd. & Najmi, A. K. Effect of statins on amyloidosis in the rodent models of Alzheimer’s disease: Evidence from the preclinical meta-analysis. Brain Res. 1749, 147115 (2020).

64. Tong, X.-K., Royea, J. & Hamel, E. Simvastatin rescues memory and granule cell maturation through the Wnt/β-catenin signaling pathway in a mouse model of Alzheimer’s disease. Cell Death Dis. 13, 325 (2022).

65. Tong, X.-K., Lecrux, C., Rosa-Neto, P. & Hamel, E. Age-Dependent Rescue by Simvastatin of Alzheimer’s Disease Cerebrovascular and Memory Deficits. J. Neurosci. 32, 4705–4715 (2012).

66. Liu, C.-C., Liu, C.-C., Kanekiyo, T., Xu, H. & Bu, G. Apolipoprotein E and Alzheimer disease: risk, mechanisms and therapy. Nat. Rev. Neurol. 9, 106–118 (2013).

67. Cai, R. et al. Role of RhoA/ROCK signaling in Alzheimer’s disease. Behav. Brain Res. 414, 113481 (2021).

68. Schmidt, S. I., Blaabjerg, M., Freude, K. & Meyer, M. RhoA Signaling in Neurodegenerative Diseases. Cells 11, 1520 (2022).

69. Oakley, H. et al. Intraneuronal beta-amyloid aggregates, neurodegeneration, and neuron loss in transgenic mice with five familial Alzheimer’s disease mutations: potential factors in amyloid plaque formation. J Neurosci 26, 10129–10140 (2006).

70. Lee, K. F. et al. Targeted Mutation of the Gene Encoding the Low Affinity Ngf Receptor P75 Leads to Deficits in the Peripheral Sensory Nervous-System. Cell 69, 737–749 (1992).

