## Supplementary material for "Neuroprotective function of astrocyte p75^NTR^ in Alzheimer’s Disease through regulation of cholesterol metabolism": S legends

### Supplementary figure legends

**Supplementary Figure S1. Generation of conditional knock-in p75$^{NTR}$ alleles ΔDD and C259A**

A) Schematic of the *p75ntr* locus (not at scale) with strategy for generation of the conditional ΔDD mutant knock-in (KI) allele.

B) Schematic of the *p75ntr* locus (not at scale) with strategy for generation of the conditional C259A mutant knock-in (KI) allele.

**Supplementary Figure S2. Expression of ΔDD$^{fl/fl}$ and C259A$^{fl/fl}$ alleles before and after Cre-mediated recombination**

(A) Expression of wild type and ΔDD$^{fl/fl}$ p75$^{NTR}$ alleles in cultured hippocampal and cortical astrocytes detected with antibodies against the receptor extracellular domain (ECD). Immunoblot with GAPDH is shown below as loading control. Three independent cultures are shown in each case.

(B) Expression of wild type and C259A$^{fl/fl}$ p75$^{NTR}$ alleles in cultured hippocampal and cortical astrocytes detected with antibodies against the receptor extracellular domain (ECD). Immunoblot with GAPDH is shown below as loading control. Three independent cultures are shown in each case.

(C) Expression of ΔDD$^{fl/fl}$ p75$^{NTR}$ allele in cultured hippocampal and cortical astrocytes after transduction with control or Cre-expressing adenoviruses (ADM-CMV-Cre) detected with antibodies against the receptor extracellular domain (ECD) or death domain (DD). Note the reduction in DD immunoreactivity in astrocytes infected with ADM-CMV-Cre viruses indicating Cre-mediated DD deletion. Immunoblot with GAPDH is shown below as loading control. Three independent cultures are shown in each case.

(D) Expression of C259A p75$^{NTR}$ allele in cultured hippocampal and cortical astrocytes after transduction with control or Cre-expressing adenoviruses (ADM-CMV-Cre) detected with antibodies against the receptor extracellular domain (ECD). Immunoblot with GAPDH is shown below as loading control. Three independent cultures are shown in each case.

**Supplementary Figure S3. Upregulation of p75$^{NTR}$ expression in astrocytes of 5xFAD hippocampus**

(A) Representative micrographs of $p75^{NTR}$ and GFAP immunohistochemistry in the hippocampus of 9 month old wild type and 5xFAD mice showing increased $p75^{NTR}$ expression in astrocytes of 5xFAD mice.

(B) Quantification of GFAP/$p75^{NTR}$ double positive cells expressed as % of all $GFAP^{+}$ cells in hippocampus of 9 month old wild type and 5xFAD mice. Results are presented as mean ± SEM (N=5 per group) and analyzed by Student's t test. ****, p<0.0001 vs. 5xFAD.

**Supplementary Figure S4. Gene sets enriched in $p75^{\Delta DD}$ and $p75^{C259A}$ hippocampal and cortical astrocytes**

Shown are top gene sets (from Reactome and KEGG pathway databases) enriched in cultured $p75^{\Delta DD}$ and $p75^{C259A}$ hippocampal and cortical astrocytes revealed by bulk RNA-Seq. Cholesterol and steroid biosynthesis pathways are highlighted.

**Supplementary Figure S5. Astrocyte expression of signaling-deficient p75NTR variants up-regulates astrocyte cholesterol biosynthesis and cholesterol content in cortical neurons**

(A) Representative micrographs showing Filipin staining and GFAP immunostaining of sagittal sections through the cerebral cortex of wild type, 5xFAD, Aldh1l1-$CreER^{T2}$;$\triangle DD^{fl/fl}$;5xFAD and Aldh1l1-$CreER^{T2}$;$C259A^{fl/fl}$;5xFAD mice at 6 month of age. Mice were injected with tamoxifen at 2 month of age. Histogram shows quantification of Filipin area (as % of GFAP area) in hippocampus of wild type, 5xFAD, Aldh1l1-$CreER^{T2}$;$\triangle DD^{fl/fl}$;5xFAD and Aldh1l1-$CreER^{T2}$;$C259A^{fl/fl}$;5xFAD mice injected with TMX at 2 month and analyzed at 6 month. Results are presented as mean ± SEM (N=7-11 mice per group) and analyzed by one-way ANOVA followed by Tukey's multiple comparisons test. *, p<0.05; **, p<0.01 vs. 5xFAD.

(B) Representative micrographs of cultured astrocytes isolated from the cortex of wild type, $p75^{\Delta DD}$ and $p75^{C259A}$ mice stained with Filipin to reveal cholesterol content and anti-GFAP antibody. Histogram shows quantification of Filipin area normalized to $GFAP^{+}$ cell number and expressed as percentage of wild type (WT) levels. Results are presented as mean ± SEM (N=3 independent experiments each performed in triplicate) and analyzed by one-way ANOVA followed by Tukey's multiple comparisons test. **, p<0.01 vs. WT.

(C) Representative micrographs of cultured wild type cortical astrocytes treated with different neurotrophins at the indicated concentrations stained with Filipin to reveal cholesterol content and anti-GFAP antibody. Histogram shows quantification of Filipin area normalized to $GFAP^{+}$ cell number and expressed as percentage of vehicle levels. Results are presented as mean ± SEM (N=3 independent experiments each performed in triplicate) and analyzed by one-way ANOVA followed by Tukey's multiple comparisons test. **, $p<0.01$; ***, $p<0.001$; ****, $p<0.0001$ vs. vehicle.

(D) Quantification of free cholesterol in the supernatant of cultured cortical astrocytes isolated from wild type, $p75^{\Delta DD}$ and $p75^{C259A}$ mice. Results are presented as mean ± SEM (N=3 independent experiments each performed in triplicate) and analyzed by one-way ANOVA followed by Tukey's multiple comparisons test. *, $p<0.05$; **, $p<0.01$ vs. WT.

(E) Representative micrographs of cultured wild type cortical neurons supplemented with conditioned medium from wild type, $p75^{\Delta DD}$ and $p75^{C259A}$ astrocytes stained with Filipin to reveal cholesterol content and anti-MAP2 antibody. Control indicates normal unsupplemented neuron culture medium. Histogram shows quantification of Filipin area normalized to MAP2 area and expressed as percentage of wild type (WT) levels. Results are presented as mean ± SEM (N=3 independent experiments each performed in triplicate) and analyzed by one-way ANOVA followed by Tukey's multiple comparisons test. ****, $p<0.0001$ vs. control.

(F) Representative micrographs of cultured astrocytes isolated from the cerebral cortex of wild type, $p75^{DHEA}$ and $p75^{KKEA}$ mice stained with Filipin to reveal cholesterol content and anti-GFAP antibody. Histogram shows quantification of Filipin area normalized to $GFAP^+$ cell number and expressed as percentage of wild type (WT) levels. Results are presented as mean ± SEM (N=3 independent experiments each performed in triplicate) and analyzed by one-way ANOVA followed by Tukey's multiple comparisons test. **, $p<0.01$ vs. WT.

(G) Representative micrographs of cultured wild type cortical astrocytes treated with NF-kB inhibitor JSH23 (20μM) stained with Filipin to reveal cholesterol content and anti-GFAP antibody. Histogram shows quantification of Filipin area normalized to $GFAP^+$ cell number and expressed as percentage of vehicle levels. Results are presented as mean ± SEM (N=3 independent experiments each performed in triplicate) and analyzed by Student's t-test. ***, $p<0.001$ vs. vehicle.

**Supplementary Figure S6. Neurotrophins have no effect on the cholesterol content of ∆DD and C259A astrocytes**

Histograms shows quantification of Filipin area normalized to $GFAP^+$ cell number and expressed as percentage of vehicle levels in hippocampal and cortical astrocytes derived from $p75^{\Delta DD}$ and $p75^{C259A}$ mice. Results are presented as mean ± SEM (N=3 independent experiments each performed in triplicate) and analyzed by one-way ANOVA followed by Tukey's multiple comparisons test.

**Supplementary Figure S7. Generation of knock-in $p75^{NTR}$ alleles DHEA and KKEA**

A) Schematic of the *p75ntr* locus (not at scale) with strategy for generation of the DHEA mutant knock-in (KI) allele.

B) Schematic of the *p75ntr* locus (not at scale) with strategy for generation of the KKEA mutant knock-in (KI) allele.

**Supplementary Figure S8. Statin treatment decreases brain cholesterol and counteracts the effects of impaired astrocyte $p75^{NTR}$ function on AD neuropathology**

(A) Representative micrographs showing Filipin staining and GFAP immunostaining of sagittal sections through the hippocampus and cortex of wild type, 5xFAD and Aldh1l1-$CreER^{T2}$;△$DD^{fl/fl}$;5xFAD mice treated with vehicle (control) or statins (companion images to Figure 8).

(B) Representative micrographs showing Aβ plaque depositions (as revealed by immunohistochemistry with 6E10 antibody) in sagittal sections through the hippocampus, cerebral cortex and thalamus of 5xFAD and Aldh1l1-$CreER^{T2}$;△$DD^{fl/fl}$;5xFAD mice treated with vehicle (control) or statins (companion images to Figure 8).

(C) Representative micrographs showing GFAP immunostaining of sagittal sections through the hippocampus, cerebral cortex and thalamus of 5xFAD and Aldh1l1-$CreER^{T2}$;△$DD^{fl/fl}$;5xFAD mice treated with vehicle (control) or statins (companion images to Figure 8).

(D) Representative micrographs showing Iba1 immunostaining of sagittal sections through the hippocampus, cerebral cortex and thalamus of 5xFAD and Aldh1l1-$CreER^{T2}$;△$DD^{fl/fl}$;5xFAD mice treated with vehicle (control) or statins (companion images to Figure 8).
