## Supplementary figures and images for "Neuroprotective function of astrocyte p75^NTR^ in Alzheimer’s Disease through regulation of cholesterol metabolism"

### s1

Supplementary. Fig. S1

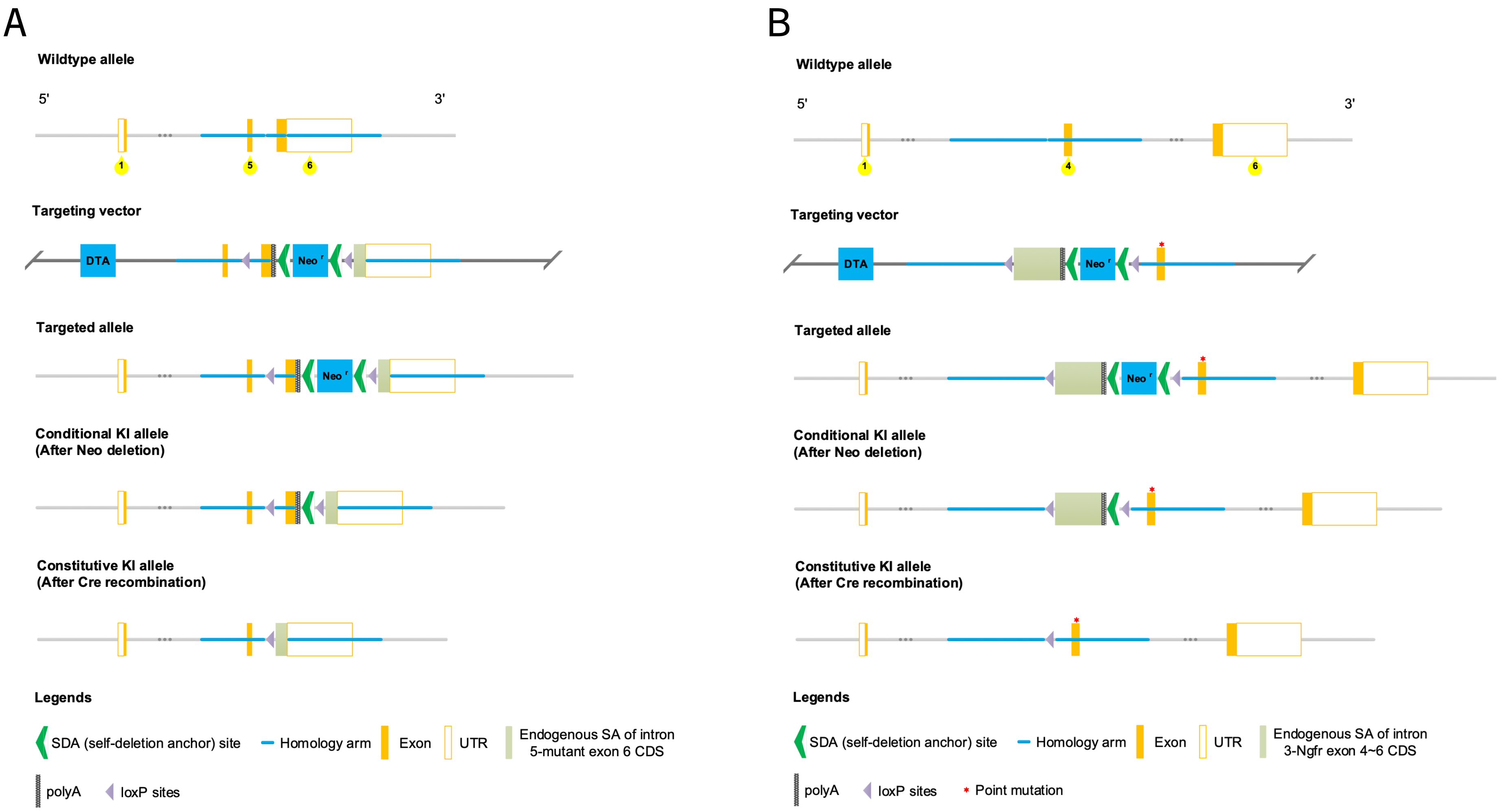

### s2

Supplementary Fig. S2

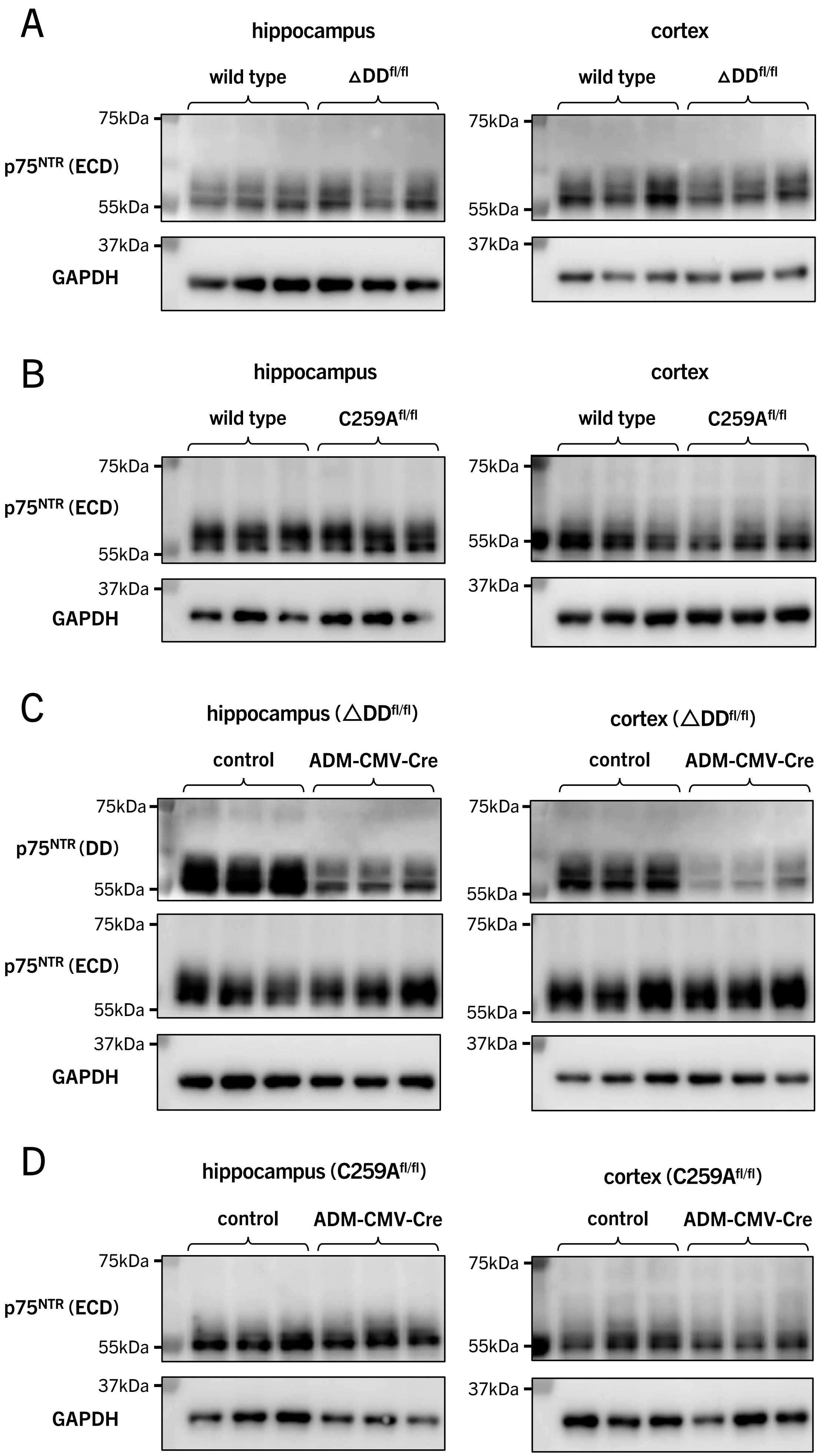

### s3

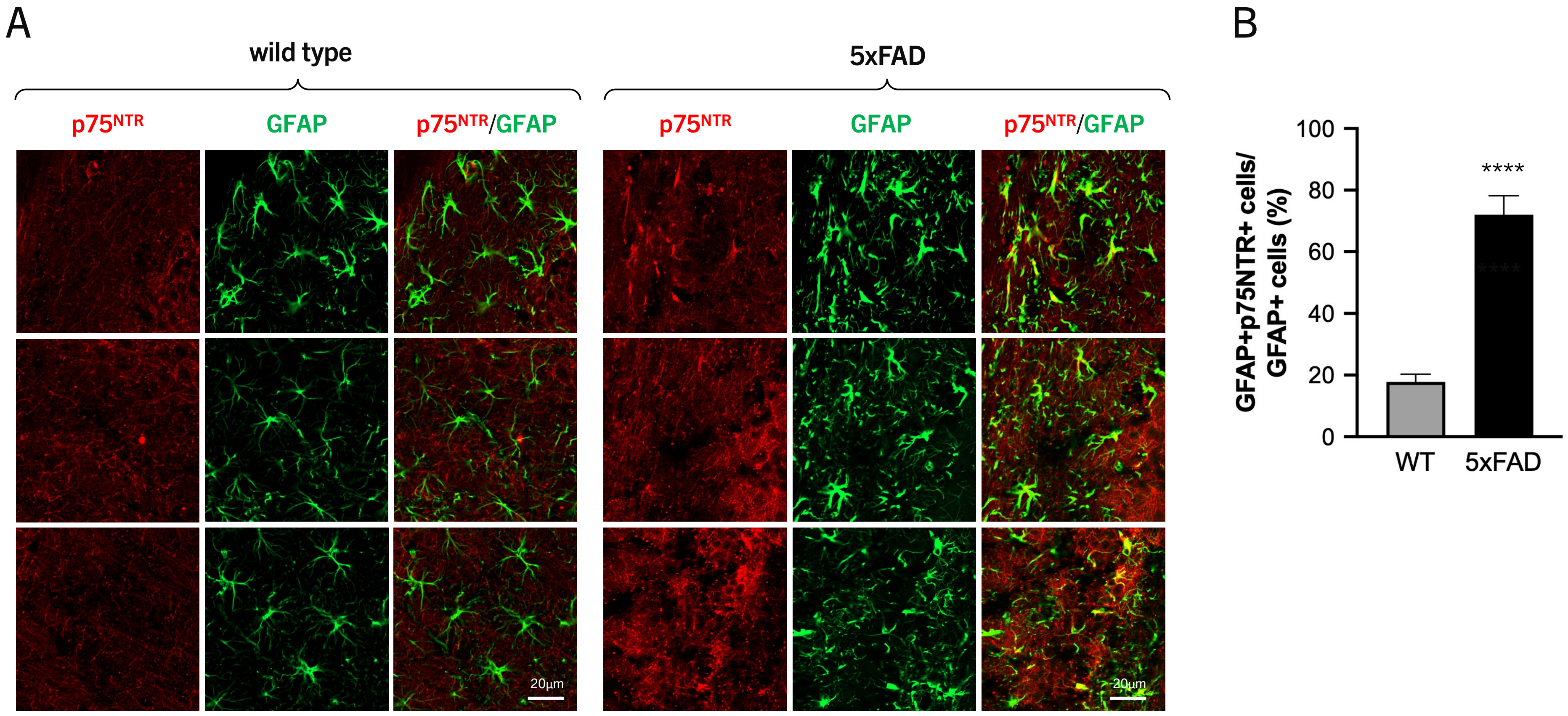

### s5

Supplementary Fig. S5

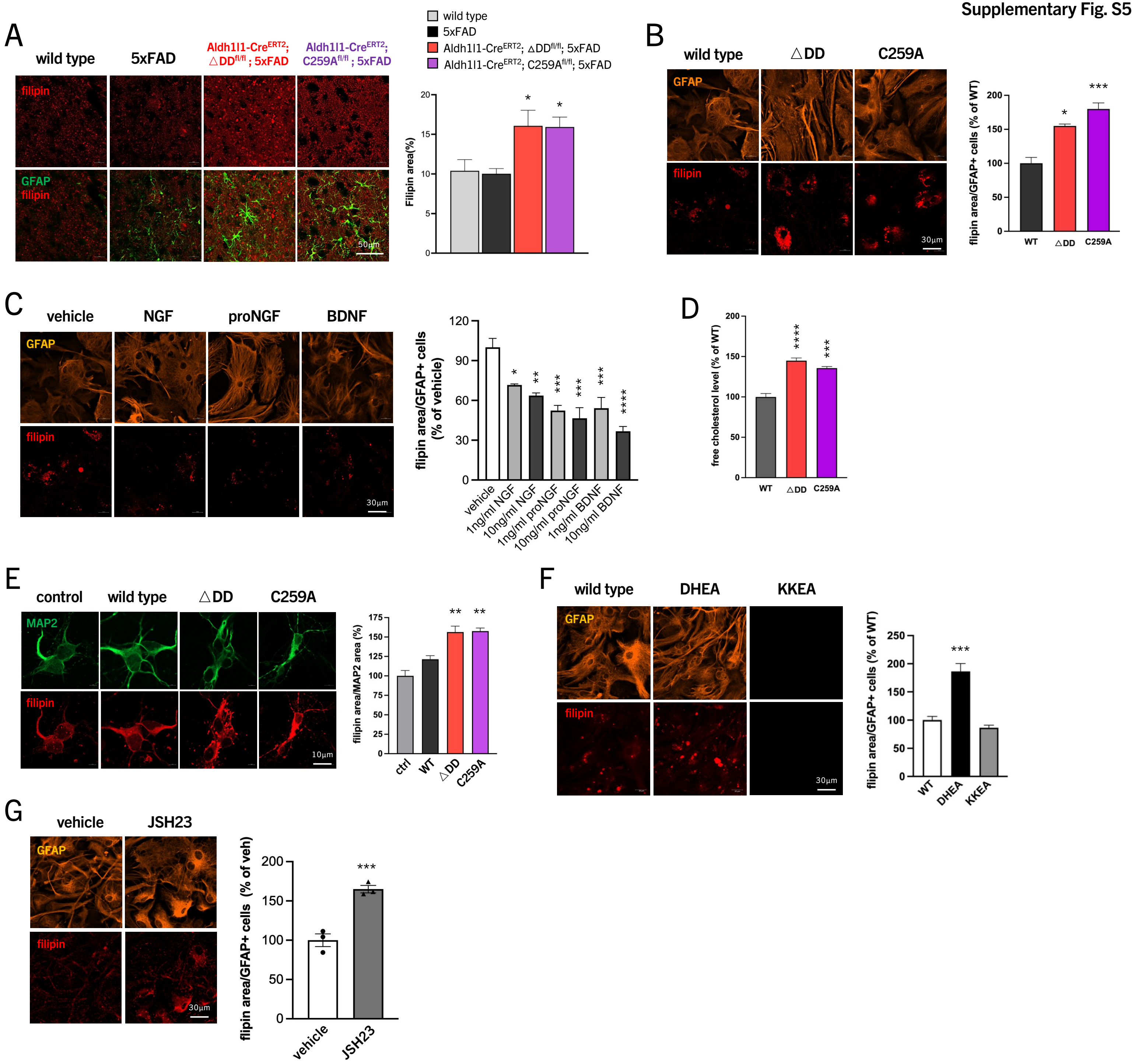

### s6

**Supplementary Fig. S6**

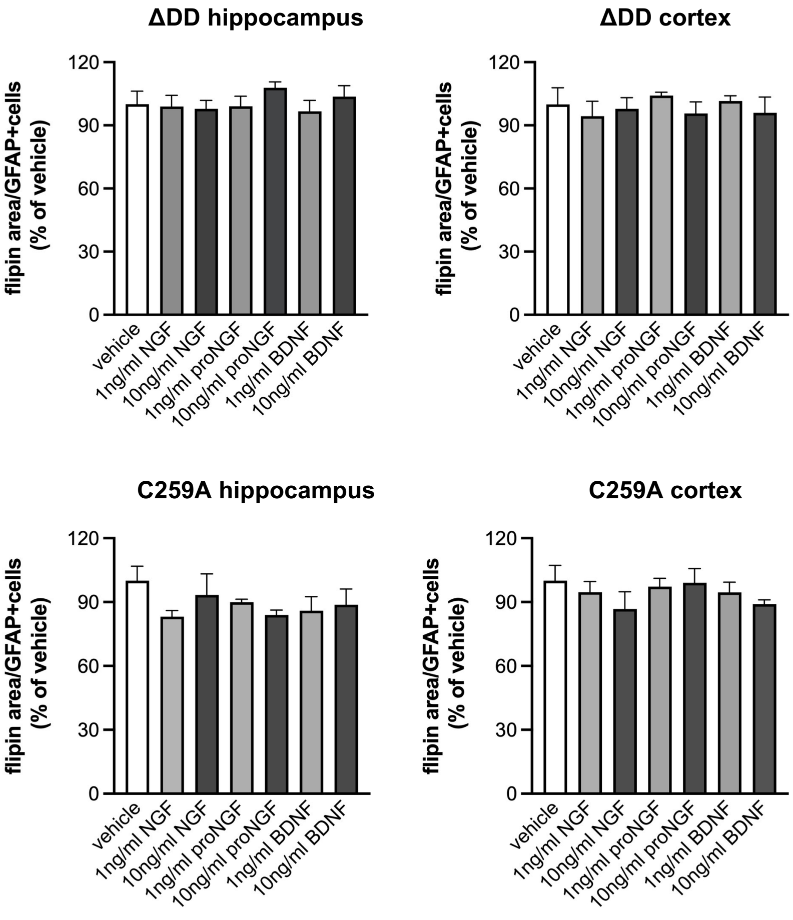

### s7

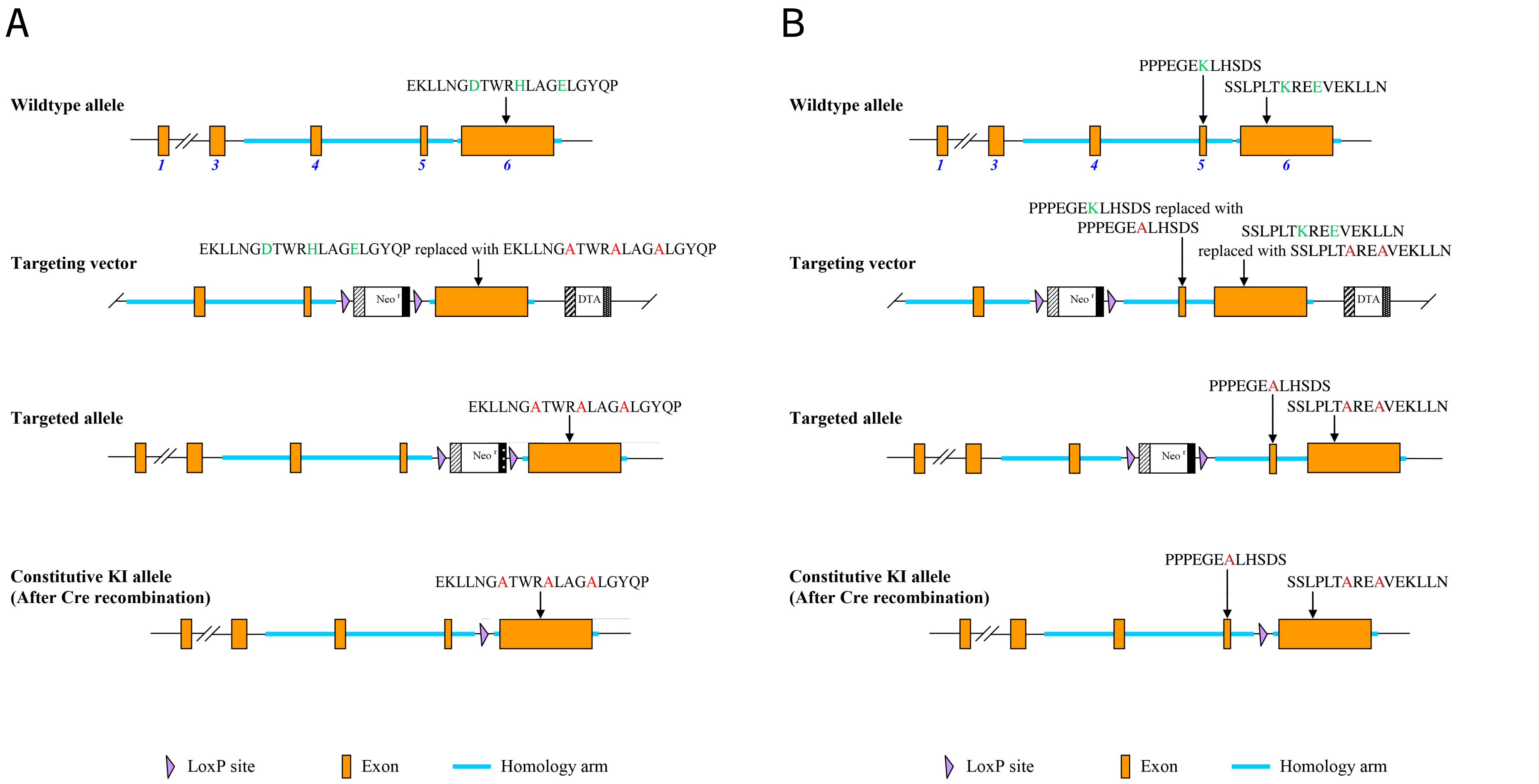

### s8

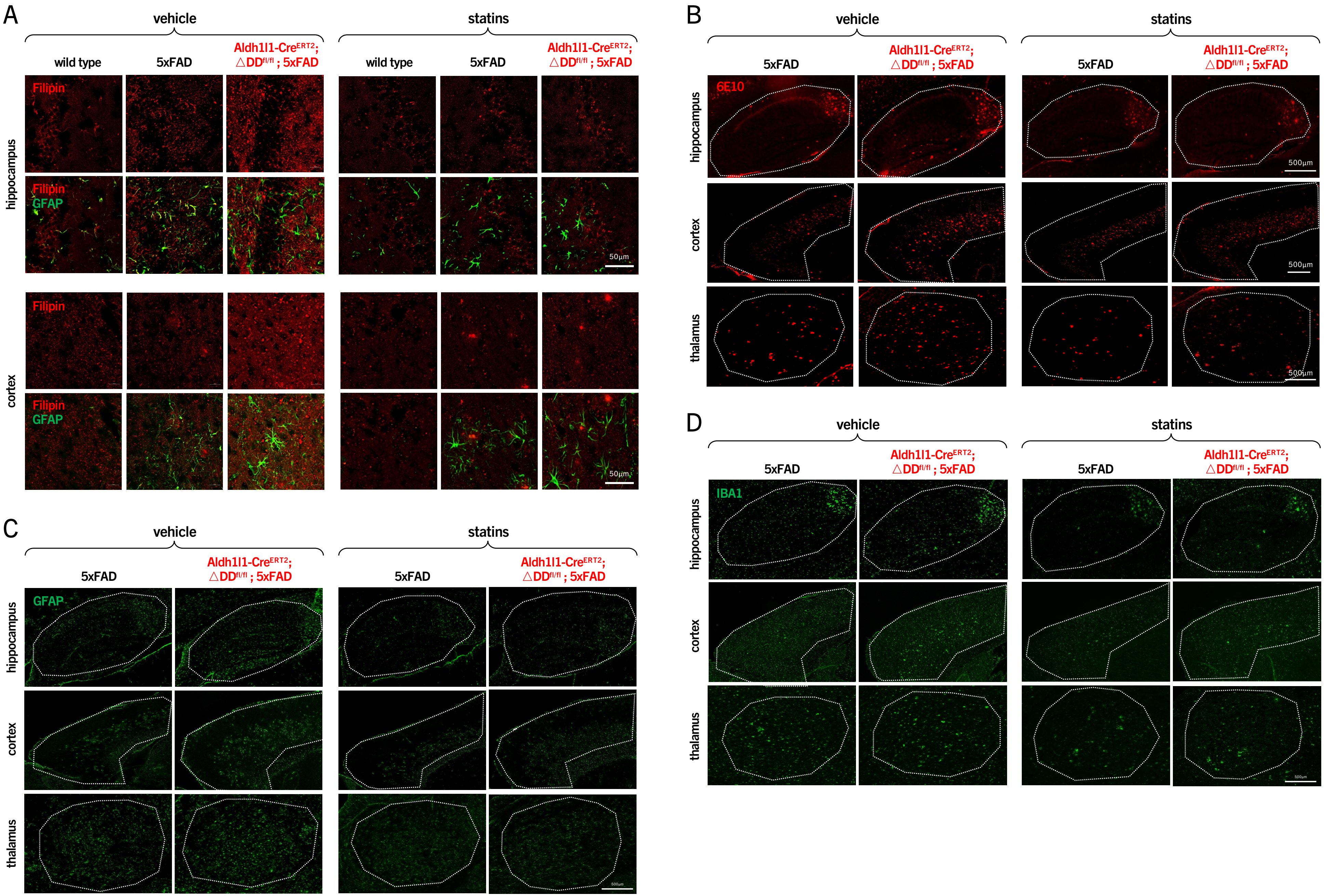
