## Supplementary material for "Neuroprotective function of astrocyte p75^NTR^ in Alzheimer’s Disease through regulation of cholesterol metabolism": s4

### Gene sets enriched in p75$^{C259A}$ cortical astrocytes

| | |
|---|---|
| 1 | CHOLESTEROL BIOSYNTHESIS(R_MMU_191273) |
| 2 | ACTIVATION OF THE PRE_REPLICATIVE COMPLEX(R_MMU_68962) |
| 3 | TELOMERE C_STRAND (LAGGING STRAND) SYNTHESIS(R_MMU_174417) |
| 4 | PROCESSIVE SYNTHESIS ON THE LAGGING STRAND(R_MMU_69183) |
| 5 | EXTENSION OF TELOMERES(R_MMU_180786) |

### Gene sets enriched in p75$^{ΔDD}$ hippocampal astrocytes

| | |
|---|---|
| 1 | LYSINE_DEGRADATION(MMU00310) |
| 2 | STEROID_BIOSYNTHESIS(MMU00100) |
| 3 | ECM_RECEPTOR_INTERACTION(MMU04512) |
| 4 | ARRHYTHMOGENIC_RIGHT_VENTRICULAR_CARDIOMYOPATHY(MMU05412) |
| 5 | ALPHA_LINOLENIC_ACID_METABOLISM(MMU00592) |

### Gene sets enriched in p75$^{C259A}$ hippocampus

| | |
|---|---|
| 1 | GLYCOSPHINGOLIPID_BIOSYNTHESIS___GANGLIO_SERIES(MMU00604) |
| 2 | HOMOLOGOUS_RECOMBINATION(MMU03440) |
| 3 | MUCIN_TYPE_O_GLYCAN_BIOSYNTHESIS(MMU00512) |
| 4 | ONE_CARBON_POOL_BY_FOLATE(MMU00670) |
| 5 | CARBON_METABOLISM(MMU01200) |
| 6 | VARIOUS_TYPES_OF_N_GLYCAN_BIOSYNTHESIS(MMU00513) |
| 7 | TYPE_II_DIABETES_MELLITUS(MMU04930) |
| 8 | CYSTEINE_AND_METHIONINE_METABOLISM(MMU00270) |
| 9 | PROTEIN_EXPORT(MMU03060) |
| 10 | DNA_REPLICATION(MMU03030) |
| 11 | PENTOSE_PHOSPHATE_PATHWAY(MMU00030) |
| 12 | MISMATCH_REPAIR(MMU03430) |
| 13 | FRUCTOSE_AND_MANNOSE_METABOLISM(MMU00051) |
| 14 | STEROID_BIOSYNTHESIS(MMU00100) |

### Gene sets enriched in p75$^{ΔDD}$ cortex

| | |
|---|---|
| 1 | CHROMOSOME MAINTENANCE(R_MMU_73886) |
| 2 | NUCLEOSOME ASSEMBLY(R_MMU_774815) |
| 3 | DEPOSITION OF NEW CENPA_CONTAINING NUCLEOSOMES AT THE CENTROMERE(R_MMU_606279) |
| 4 | G2_M DNA DAMAGE CHECKPOINT(R_MMU_69473) |
| 5 | SUMOYLATION OF CHROMATIN ORGANIZATION PROTEINS(R_MMU_4551638) |
| 6 | TRANSCRIPTIONAL REGULATION BY SMALL RNAS(R_MMU_5578749) |
| 7 | DNA DAMAGE_TELOMERE STRESS INDUCED SENESCENCE(R_MMU_2559586) |
| 8 | ACTIVATION OF ATR IN RESPONSE TO REPLICATION STRESS(R_MMU_176187) |
| ⋮ | |
| 29 | TNFR2 NON_CANONICAL NF_KB PATHWAY(R_MMU_5668541) |
| 30 | DNA DOUBLE_STRAND BREAK REPAIR(R_MMU_5693532) |
| 31 | CHOLESTEROL BIOSYNTHESIS(R_MMU_191273) |
| 32 | SIGNALING BY NON_RECEPTOR TYROSINE KINASES(R_MMU_9006927) |
| 33 | SIGNALING BY PTK6(R_MMU_8848021) |
